# A Survey of the *Desulfuromonadia* cytochromome provides a glimpse of the unexplored diversity of multiheme cytochromes in nature

**DOI:** 10.1101/2024.01.05.574438

**Authors:** Ricardo Soares, Bruno M. Fonseca, Benjamin W. Nash, Catarina M. Paquete, Ricardo O. Louro

## Abstract

Multiheme cytochromes (MHC) provide prokaryotes with a broad metabolic versatility that contributes to their role in the biogeochemical cycling of the elements. However, MHC were isolated and studied in detail only from a limited number of species. To obtain a broader view of the diversity of MHC, we employed bioinformatics tools to study the cytochromome encoded in the genomes of the *Desulfuromonadia* class. We found that MHC predicted to be extracellular are the least conserved and present higher diversity. Although the most prevalent MHC have homologues already characterized, nearly half of the MHC families in the *Desulforomonadia* class have no known homologues and AlphaFold2 was employed to predict their 3D structures. This work illuminates for the first time the universe of experimentally uncharacterized cytochromes that are likely to contribute to the metabolic versatility and to the fitness of *Desulfuromonadia* in diverse environmental conditions and to drive biotechnological applications.

## Introduction

Multiheme *c*-type cytochromes (MHC) are metalloproteins found widespread across the Bacteria and Archaea domains that play central roles in diverse anabolic and catabolic pathways. These pathways contribute significantly to the biogeochemical cycling of key elements such as metals (e.g. iron), nitrogen, and sulfur (*1*, *2*). The functional versatility of these proteins reflects their tunable coordination, redox, spin, and acid–base properties. MHC contain two or more heme cofactors covalently bound to the protein polypeptide chain. This typically occurs through two thioether bonds between two nearby cysteines of the apoprotein and the two vinyl groups of the heme cofactors (*3*). Although the most common heme-binding motif is CX_2_CH, where ‘X’ represents any possible amino acid, other less common heme-binding motifs are described in the literature, namely CXCK (*4*), CX_2_CK (*5*), CX_3_CH (*6*), CX_4_CH (*7*), CX_11_CH, CX_15_CH (*8*) and CX_17_CH (*9*). In all these cases, the iron is coordinated by the nitrogen atoms of the protoporphyrin plane and is axially coordinated by residues of the protein, forming a square pyramidal (penta-coordinated) or octahedral coordination (hexa–coordinated) geometries. The proximal axial ligand comes from the histidine or lysine that follows the second bound cysteine in the polypeptide chain (*2*).

The known oligomerization states of MHC range from monomers to the current maximum of an octamer of trimers (24-mer) for the Hydrazine Dehydrogenase complex, containing a total of 192 hemes that are closely packed to facilitate fast electron transfer (*10*). Another oligomerization mode is found in the so-called extracellular cytochrome nanowires from electroactive bacteria (*11–13*) and archaea (*14*). These proteins are composed of repetitive MHC subunits that extend outwards from the cell surface reaching micrometer-long lengths. Concerning the number of heme cofactors per polypeptide chain, molecular structures were experimentally obtained for MHC spanning from dihemes (*15*) to the hexadecaheme HmcA (*16*). However, genome mining has revealed putative MHC with significantly higher numbers of heme cofactors per polypeptide chain. The current record holder is a sequence encoding a putative MHC containing 113 heme-binding motifs (*17*). The extensive variety of existing MHC was observed to be a result of a complex process of fusion and fission processes involving heme and protein modules across time in a widespread group of MHCs (*9*, *18–20*).

The first bioinformatics investigation of the rich diversity of MHC as a whole analyzed 594 representative prokaryotic genomes (*21*). From these genomes, 258 were found to contain sequences coding for MHC, resulting in the identification of a total of 1659 MHC sequences. This analysis revealed a significant gap in our understanding of the ‘cytochromome’. Out of the 124 clusters generated from sequence alignments, only 12 had representative structures, accounting for approximately 10 % of the total clusters. Since then, many MHC were characterized (*2*, *10*) and among prokaryotes, the *Desulfuromadia* class (former *Desulforomonadales* class (*22*)) comprises members with the highest known number of predicted MHC per genome. For example, the genome of *Geotalea uraniireducens* Rf4 (former *Geobacter uranireducens* (*22*)) was reported to contain a total of 75 MHC, constituting approximately 1,7% of all the protein-coding genes in the genome (CP000698.1). This organism stands out as having the highest number of MHC per genome identified thus far (*21*). In addition, *Desulfuromadia* genomes also contain protein-coding sequences that exhibit some of the highest numbers of heme-binding motifs per MHC sequence ever found. Specifically, protein-coding sequences with 69 and 53 heme-binding motifs were identified in this group (*23*, *24*). Members of this class of organisms participate in the biogeochemical cycle of elements, particularly of metals, and perform extracellular electron transfer to exchange electrons with metallic minerals in their environment (*25*) or to electrodes where they generate some of the highest current densities ever recorded in bioelectrochemical systems (*26*). These systems use microorganisms as living catalysts to exchange electrons with electrodes to develop sustainable industrial applications, such as electricity production coupled with wastewater treatment, water desalination, biosensing and production of added-value compounds (*27*). The best-studied microorganism of the *Desulfuromadia* class is *Geobacter sulfurreducens* PCA, which together with the *Gammaproteobacterium Shewanella oneidensis* MR-1 are model electroactive organisms that contain extracellular electron transfer (EET) pathways fully composed by MHC. To a lesser extent, other *Desulfuromadia* members have also been studied in terms of current production in bioelectrochemical systems and/or fundamentals of EET. These include other *Geobacter* spp. (*28–30*), *Desulfuromonas acetoxidans* (*31–33*) and *Geoalkalibacter* spp. (*34*, *35*). In this study, we focused on the ‘cytochromome’ of the *Desulfuromadia* class and obtained a comprehensive overview of the diversity of MHC in these organisms and conservation patterns of their number of heme-binding motifs. This analysis highlighted MHC variants with predicted topology and heme content that have not been previously explored through genetic, biochemical, or structural biology methods. These findings identify significant MHC candidates for future research, particularly in the development of sustainable bioelectrochemical technologies or for the understanding of the molecular foundations of metabolic pathways sustaining key biogeochemical cycles of the elements.

## Results

### Distribution of the number of heme-binding motifs per MHC reflects conflicting evolutionary pressures

We gathered a total of 84 genomes from members of the *Desulfuromonadia* class (Data S1). These genomes are rich in MHC and yielded 4716 protein sequences identified as MHC within this dataset (Data S2). This contrasts with just 1659 MHC sequences previously identified in a much larger sample of 594 prokaryotic genomes from diverse phylogenetic backgrounds (*21*). The current result represents an almost 3-fold increase in the number of MHC studied. More importantly, it represents a nearly 20-fold increase in the average number of MHC per genome with 56.1 MHC in the present dataset vs 2.8 MHC in the previous dataset of diverse prokaryotes, reflecting the particular context of the bioenergetic electron transfer chains of *Desulfuromonadia* members. Heme-binding motifs of the 4716 MHC protein sequences were identified showing an overall trend in the shape of the distribution of frequency of heme binding motifs per polypeptide similar to that previously found (Fig. 1). Overall, the distribution of heme-binding motifs per polypeptide is not monotonic and predominantly favors small numbers of heme-binding motifs with high frequency counts at the beginning. The data show a moderate increase in prevalence from two to five heme binding sites per polypeptide followed by a fast decay until 12 heme binding sites per polypeptide, with 87% of all putative MHC found in this range. This is then followed by a long tail with very few examples which extends until 87 heme-binding sites per polypeptide. It is nonetheless interesting to observe a spike in prevalence with a maximum at 27 heme binding sites, a broad maximum centered at 35 heme binding sites and a lone spike at 56 heme binding sites (Fig. 1). Sequence alignments showed that all of these cases (27, 35 and 56) are composed by mixed populations and therefore congregate different MHC families. The spikes at 27 and 56 are composed of sequences predicted to be either periplasmic or extracellular MHC. The peak at 35 is composed solely of sequences predicted to be extracellular MHC.

**Fig. 1.**
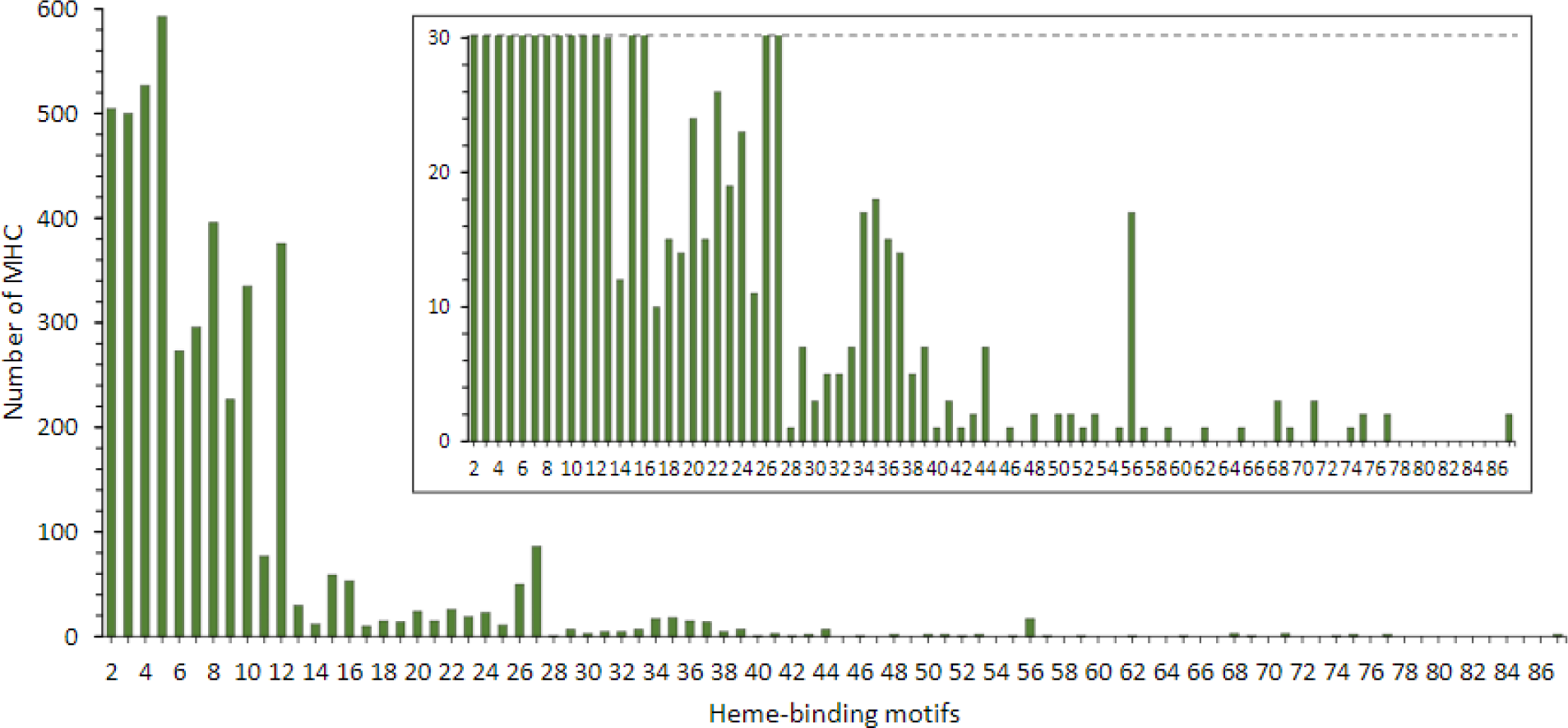
Distribution of the number of heme-binding motifs across the MHC of all the *Desulfuromonadia* genomes analyzed. Insert represents a zoomed-in view of the graph for better visualization of the small prevalences at the high numbers of putative heme-binding motifs per polypeptide.

The fact that the prevalence of MHC does not follow a monotonic function of the number of hemes likely indicates that conflicting fitness pressures operate in the development of MHC. Factors promoting a low number of hemes per MHC polypeptide chain, may include: (1) the pressure to maintain function while conserving iron and heme resources for metabolic efficiency; (2) limitations related to protein size imposed by considerations like lipid membrane thickness, periplasmic space width, and trafficking limitations for large MHC from their biogenesis in the inner-membrane of Gram-negative bacteria to their final location; (3) The fact that as the number of hemes in a MHC increases, the number of options for connecting the distal ligand of the hemes increases with a power of two, making it more difficult to ensure the correct fold of the MHC; (4) the fact that there is no obvious pressure from the side of biological catalysis to go beyond eight hemes in a MHC, given that eight electrons is the maximum necessary for reactions of biological relevance such as the conversions of sulfate to sulfide, nitrate to ammonia or CO_2_ to methane, and; (5) the option of protein polymerization over producing a single, long polypeptide chain of high molecular weight, as observed in nanowires of electroactive bacteria (*11–13*) and archaea (*14*). By contrast, the long tails may reflect specific requirements, of which two can be envisaged on the bases of current knowledge: (1) storage of electrons (*36*), and; (2) long-distance electron transfer (*13*). The combination of these pressures resulted in the most common number of heme-binding motifs per sequence in the *Desulfuromonadia* MHC being five, followed by four, two, three and eight. This result is slightly different from that reported earlier (*21*) where it was found that four was the most common number of heme-binding motifs, followed by two, five, ten and eight. These differences can be rationalized on two accounts: (1) (*21*) analyzed prokaryotes in general, while we focused on the *Desulfuromonadia*, which along with being more MHC-rich than the average of known prokaryotes, also seem to have differences in the distribution of heme-binding motifs; (2) (*21*) only considered the canonical CXXCH heme-binding motif, while we also take in consideration the other non-canonical heme-biding motifs. For example, the pentaheme NrfA, has four hemes with the canonical heme-binding motifs and frequently one catalytic heme with the CXXCK binding site (*5*). As NrfA is highly ubiquitous in prokaryotes (*9*), this likely contributed to underestimate heme-binding motif counts, skewing the count of the most common number of heme-binding motifs from five to four in (*21*). The example of NrfA hints at the balance of the conflicting pressures likely to be responsible for limiting the number of heme-binding motifs in MHC. Indeed, the five hemes of NrfA cannot hold all the necessary electrons to perform the six-electron reduction of nitrite to ammonia (*5*) However, natural selection found it more efficient to make the protein operate as a dimer than adding the additional necessary heme. Indeed, oligomerization may be a strong driver of the maximum at 5 heme-binding motifs per MHC since these are also present in homologues of the families OmcS and OmcT (next section, Fig. 4) that are highly oligomerized as cytochrome nanowires and also abundant in our data set (*13*).

### MHC repertoire shows differences between Geobacteriales and Desulfuromonadales

Our data show that the *Desulfuromonadia* class is more MHC-rich than the average found in a broad sample of prokaryotes (*21*). They also show that members of the *Geobacterales* order typically presented a higher number of MHC per genome than members of the *Desulfuromonadales* order with an average of 64 and 39, respectively (Fig. 2). In line with this observation, the elevated iron content of *G. sulfurreducens*, that constitutes 0.2 % of the cell dry weight, was proposed to result from the presence of MHC (*37*). *Geotalea uranireducens* displayed the highest number of MHC (90 MHC) corresponding to 2 % of all the genes encoded in its genome. By contrast, *Geobacter* sp. DSM 2909 presented an unexpectedly low number of MHC for a strain affiliated with the *Geobacter* genus. For the *Desulfuromonadales* order, *Desulfuromonas versatilis* presented the highest number of MHC, with a total of 80. This observation is in line with the fact that *Desulfuromonas versatilis* exhibits enhanced anaerobic respiratory versatility compared to other species of the *Desulfuromonadales* order (*24*). On the other hand, members of the *Syntrophotalea* genus are usually described as fermentative (*38*), which agrees well with the fact that for species of this genus, we could find little to no MHC. This contrasts with our discovery of 62 putative MHC in the genome of *Malonomonas rubra* DSM 5091 which was described as strictly fermentative. However, the capacity for extracellular electron transfer of this organism was not tested (*39*).

**Fig. 2.**
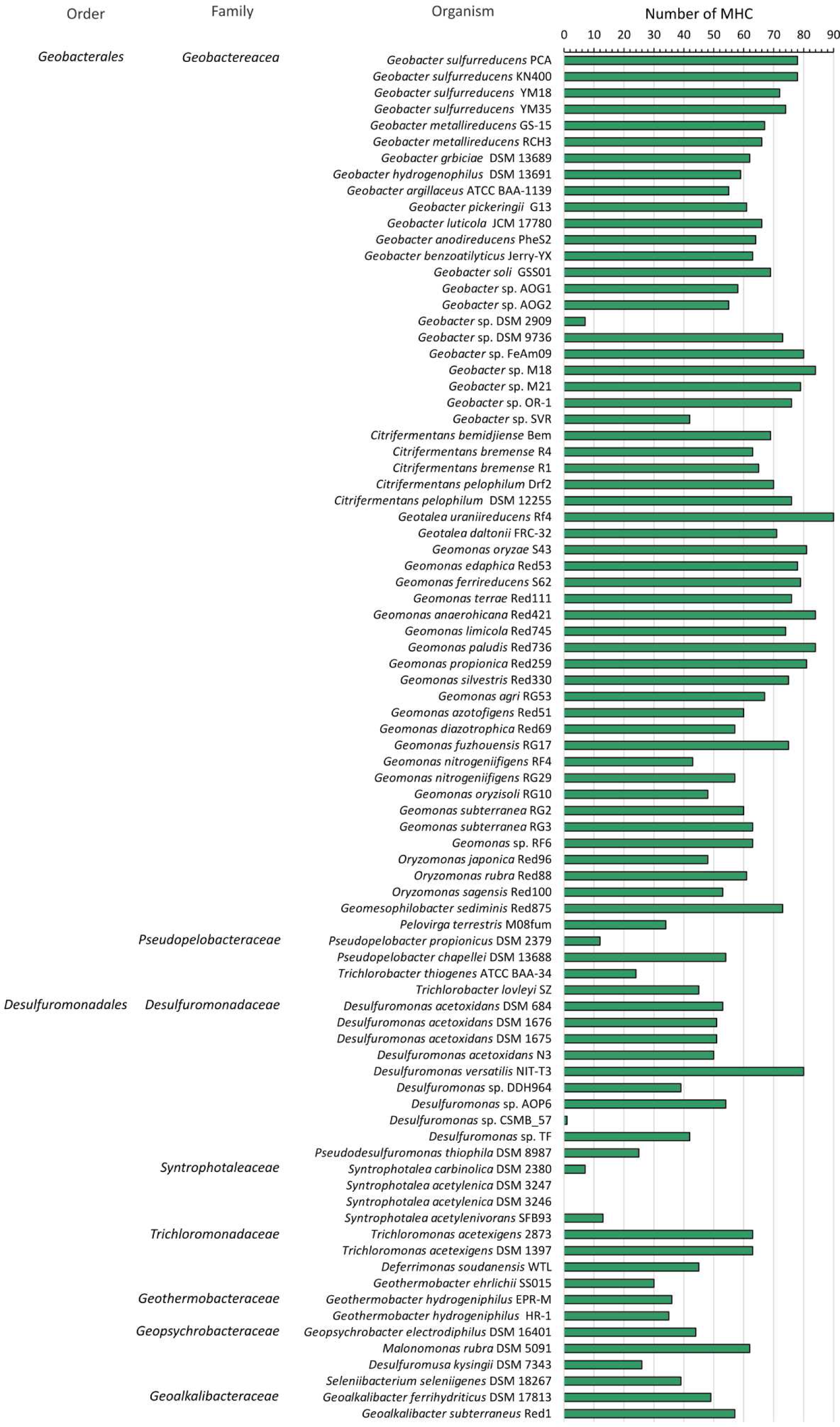
Number of putative MHC coding sequences per genome. MHC counts for each *Desulfuromonadia* genome are shown. Taxonomical classification at the level of family and order is presented on the left-hand side.

### MHC with a large number of hemes are typically outliers within the genome

Heme-binding motif distributions in individual species follow the global trend, with the median number of heme-binding motifs in the range of 2 to 9 (Fig. 3). Some species do not exhibit a distribution characterized by a long tail of MHC with high numbers of heme-binding motifs. Examples of such species include strict fermenters, such as *Syntrophotalea acetylenivorans* (*38*) and *S. carbinolica* (*40*). In addition, some species that respire soluble and insoluble electron acceptors display distributions of heme-binding motifs that lack the long tail, such as *Trichlorobacter lovleyi* (*41*) and *T. thiogenes* (*42*). Some species, like *Geobacter* sp. DSM 9736 and *Geobacter* sp. OR-1 (*43*) and *Citrifermentans pelophilum* DSM 11255 (*44*) display a large interquartile range. In these cases, these species present a high frequency of MHC with a large number of heme-binding motifs.

**Fig. 3.**
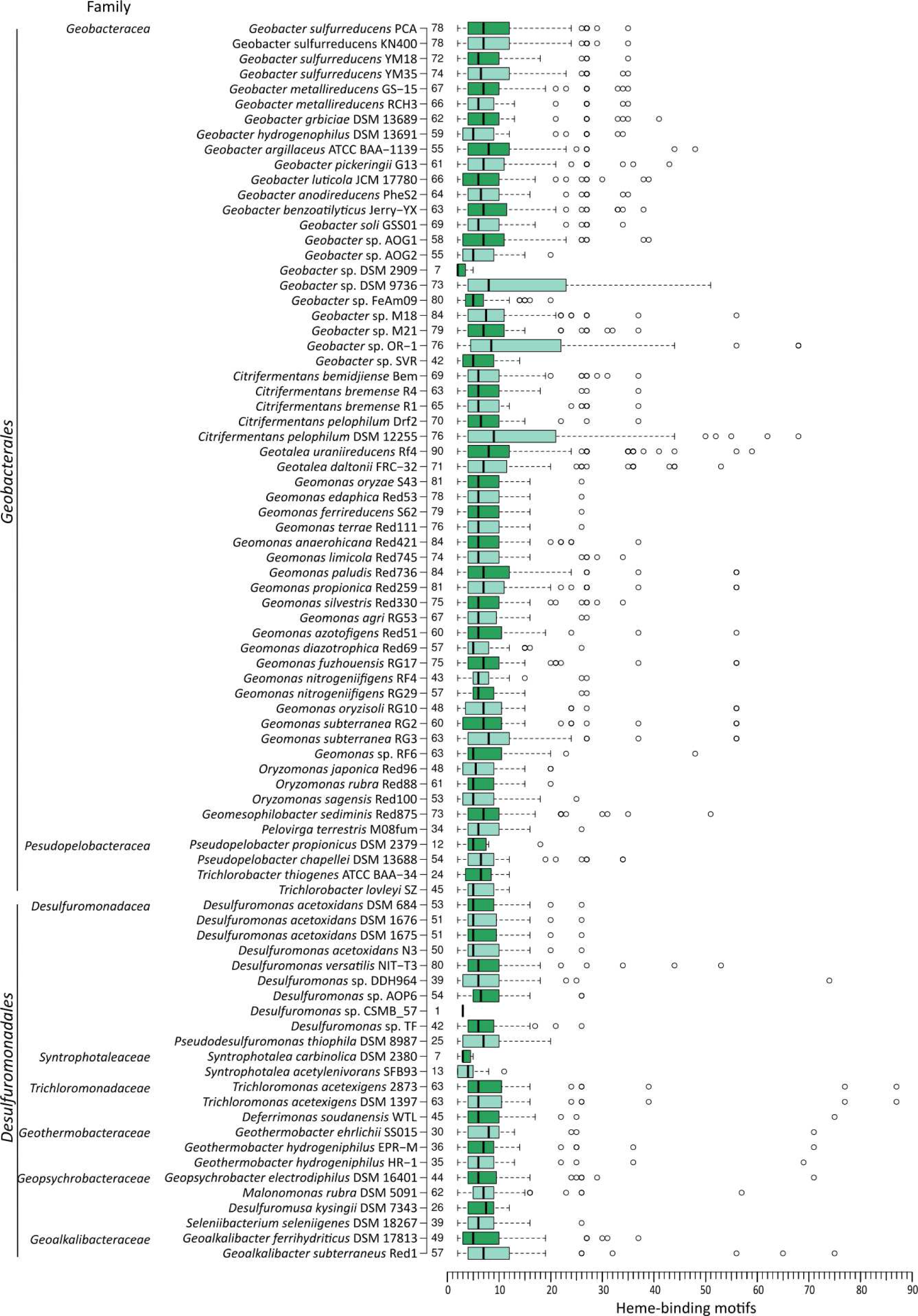
Distribution of the number of heme-binding motifs for each genome. Family and order are indicated at the left of each organism, while the number of MHC is indicated at the right. The whiskers of the boxplots extend to data points that are less than 1.5 times the interquartile range away from 1^st^/3^rd^ quartile (Turkey whiskers). Circles represent outlier values using the above criteria.

### Nearly half of the MHC in Desulforomonadia have no known homologues

To characterize the large pool of MHC from the *Desulfuromonadia*, we constructed a curated database to provide templates of MHC described in the literature that could be used to define clusters based on sequence homology (see Materials and Methods; Data S3). This database included MHC that were characterized genetically, biochemically, or structurally. Despite using template MHC that in some cases were only characterized through gene knockout studies, a significant portion of the clusters (45.5% of the dataset) remained to be identified. We were able to identify homologues for 54.4% of the *Geobacterales* MHC and 54.6% of the *Desulfuromonadales* MHC. These homologues have diverse numbers of heme binding sites as a consequence of the modular evolution of MHCs involving grafting and pruning of protein modules with heme binding sites (*9*). The clusters with the most abundant sequences are homologous to MHC already characterized (Fig. 4). For a significant number of clusters, the characterization did not include structural information and this was predicted using Alphafold2 (Fig. S1). We found homologues of putative inner membrane quinone/quinol oxidoreductases such as ActA (*45*), CbcA (*46*), CbcL (*47*), CbcS (*48*), CbcX (*48*), CymA/NapC/NrfH(*49*), and ImcH (*50*). Homologues of the periplasmic MHC were also found, including of BthA/MacA/MauG/PsaCCP (*51*), cytochrome *c*_3_ (*52*), cytochrome c_7_/PpcA/B/C/D/E (*53*), DHC2 (*54*), DsrJ (*55*), FccA (*56*), GSU0105 (*57*), GSU0616 (*58*), GSU1786 (*48*), GSU1996/OmcQ (*36*), HAO/HDH (*59*), IhOCC (*60*), MccA (*8*), NrfA (*5*), PccG(*48*), PccR (*48*), PpcF/OTR (*61*), QHNDH (*62*), sb-DHC (*63*), Split Soret cytochrome *c* (*64*), and STC (*65*). As for homologues of outer membrane MHC we found DmsE/MtoA/MtrA/OmcI (*19*), ExtA (*66*), ExtK (*66*), ExtG (*66*), and OmaC/B (*67*). Extracellular MHC were also found, including OmaV/W(ExtC) (*66*), ExtD (*66*), GSU1334/OmcZ (*11*), GSU2884(OmcA) (*48*), MtrC/F (*19*), OmcB/C (*68*), OmcE/OmcP (*12*), OmcG (*69*), OmcH/O (*69*), OmcJ/S/T (*13*), OmcM (*48*) and PgcA (*70*).

**Fig. 4.**
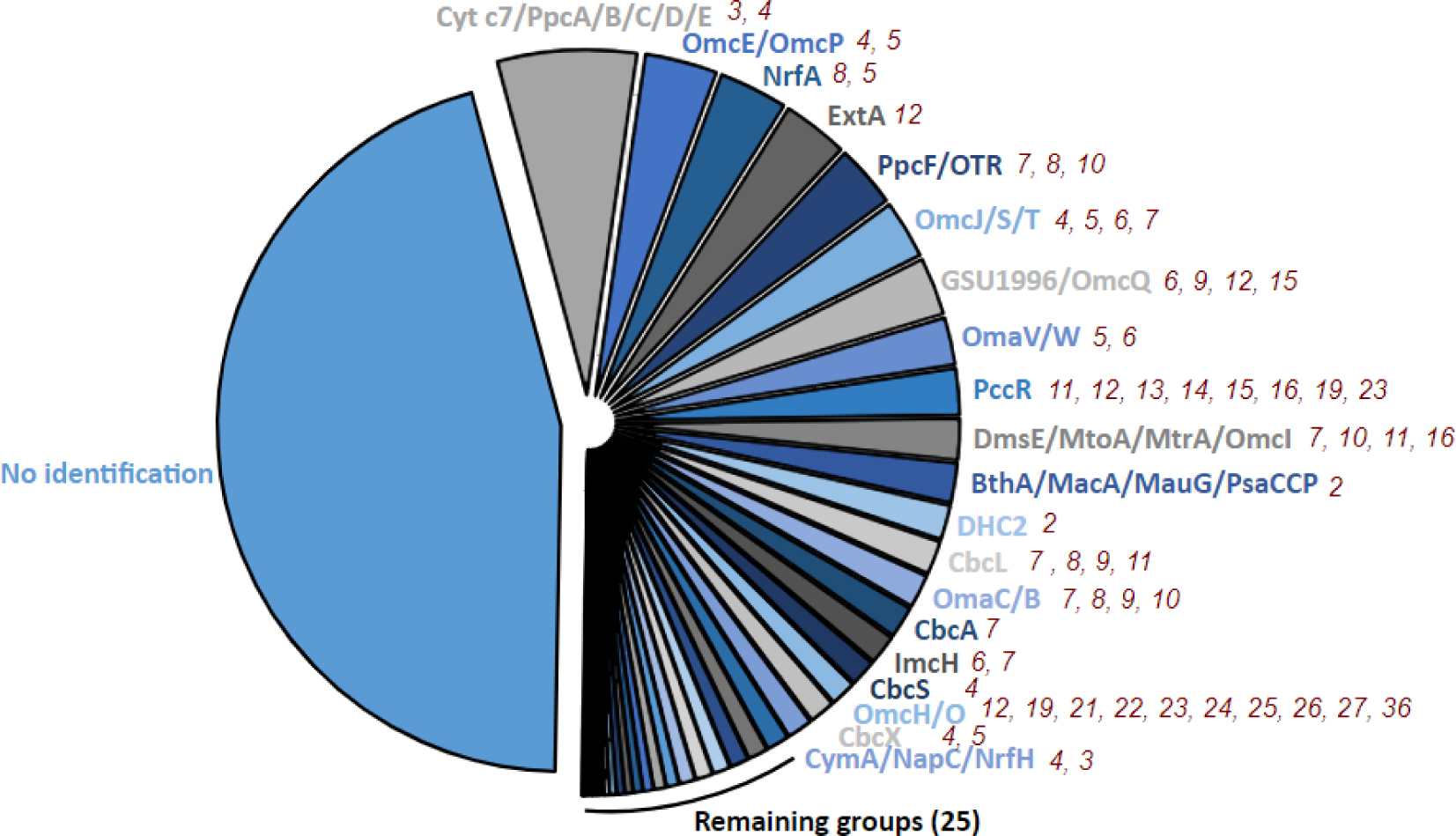
Pie-chart distribution of the number of sequences of the homologous groups identified within the MHC dataset from the. *Desulfuromonadia*. No identification represents groups that do not display significant homology to representative sequences. Out of the 45 homologous groups identified, the top 20 largest ones are labelled. The number of heme-binding motifs found in each homologous group in members of *Desulfuromonadia* is presented near each group. Detailed information can be accessed in the Table S1.

The identification process showed that the most prevalent homologous group (12.0% of the identified groups) consisted of homologs of the well-known periplasmic triheme cytochromes: *c*ytochrome *c*_7_ and PpcA-E. These are amongst the most abundant MHC found in *Desulfuromonadia* members (*32*, *71*, *72*). The second most prevalent group (6.2% of the identified groups) was the OmcE/P group. Additionally, a considerable portion of the sequences was categorized into groups comprising nanowire MHC, such as the ones that included the well-characterized OmcS (5.0%), GSU1996 (4.9%) and OmcZ (1%). This totalized 17.1% of the identified groups, which demonstrates the importance of ‘nanowire-like’ MHC for *Desulfuromonadia*. Homologous sequences for the catalytic MHC NrfA/ONR and OTR were also abundant and together constitute 11.3% of the identified sequences. Inner membrane-associated MHC seem to be less abundant overall, where CbcL homologues’ sequences were found to be dominant, representing 2.9% of the sequences that could be identified. Surprisingly, a substantial number of sequences homologous to the DmsE/MtoA/MtrA/OmcI group was also found, which were never reported in *Geobacter* or related species thus far. This group includes well-characterized proteins involved in the reduction and oxidation of soluble and insoluble compounds, organized in what is referred to as porin-cytochrome complexes that permeate the outer-membrane of several Gram-negative bacteria (*73*). A similar topological organization is also expected in members of the *Desulfuromonadia* class which show DmsE/MtoA/MtrA/OmcI in the same operon of the MtrB/PioB-like porin gene. Even more surprising, some members, in particular from the *Geobacterales* order, possess operons containing genes annotated as “ABC transporter ATP-binding protein” and “ABC transporter substrate-binding protein”, in addition to the DmsE/MtoA/MtrA/OmcI and MtrB/PioB genes. This suggests that in these organisms, DmsE/MtoA/MtrA/OmcI MHC could have a role in the transport of substances, such as the uptake of metals (*74*).

### Distribution of MHC within species shows differences between *Desulfuromonadales* and *Geobacteriales*

To relate this information with the corresponding species we represented the distribution of identified MHC by homology with each species (Fig. 5; Data S4). Within the inner membrane, ActA is the least represented in *Desulfuromonadia,* appearing in only 26% of the species. CymA/Nap/NrfH is poorly represented in the *Geobacterales*, while CbcS, CbcX and ImcH are poorly represented in the *Desulfuromonadales* (≤ 50% of the species). On the other hand, CbcA and CbcL are both well distributed among the *Desulfuromonadia* (present in ≥ 84% of the species). Typically, each genome contains only one copy of these inner membrane MHC sequences. However, there are a few exceptions to this, such as *Desulfuromonas acetoxidans* strains that have 3 or 4 CymA/Nap/NrfH genes per genome. Periplasmic MHC genes are globally dominated by the cytochrome *c*_7_/PpcA-E family. With a few exceptions, this family is widely distributed, featuring multiple paralogues per genome. In addition, BthA/MacA/MauG/PsaCCP, DHC2, NrfA, PccR and PpcF/OTR groups are also well distributed within the *Desulfuromonadia* class (≥ 54% of the species). The GSU0105 and GSU1996/OmcQ groups are widely distributed, but only within the *Geobacterales* order. MccA, QHNGH and sb-DHC were only found in one species, while cyt *c*_3_, MtrC/F and DsrJ were only found in two species. The low conservation suggests that these genes likely originated from horizontal gene transfer from other families where they are more prevalent, including *Shewanellacea* for MtrC/F (*75*) or *Desulfovibrionaceae* for cytochrome *c*_3_ (*2*). From the sequences related to MHC predicted to be located at the outer membrane, DmsE/MtoA/MtrA/OmcI, ExtA, OmaC/B are well distributed in the *Desulfuromonadia* class (≥ 58% of the species). The remaining MHC are not well distributed within the *Desulfuromonadia* class. ExtK is only well distributed in the *Geobacterales,* while ExtG is poorly distributed overall. For the sequences related to extracellular MHC the scenario is quite different since there is a pattern of global low conservation among species. The only MHC well distributed among *Desulfuromonadia* class is OmcE/P (≥ 54% of the species), while OmcH/O and OmaV/W (ExtC) are well distributed among *Geobacterales* and OmcJ/S/T group in the *Desulfuronadales* order. This suggests that exoelectrogenic EET in this class of organisms is less restrained, having more room for diversification. This result aligns with earlier bioinformatics studies conducted on the *Geobacterales* order (*76*) and *Shewanellaceae* (*75*), as well as experimental evolutionary studies on *Geobacter sulfurreducens*, focusing on adaptation to microbial fuel cells (*77*). Almost half of the sequences could not be classified and it remains uncertain whether this pattern will subsist in these cases, or if MHC families yet to be discovered are so distinct from those currently known that even the conservation pattern according to their location is significantly different.

**Fig. 5.**
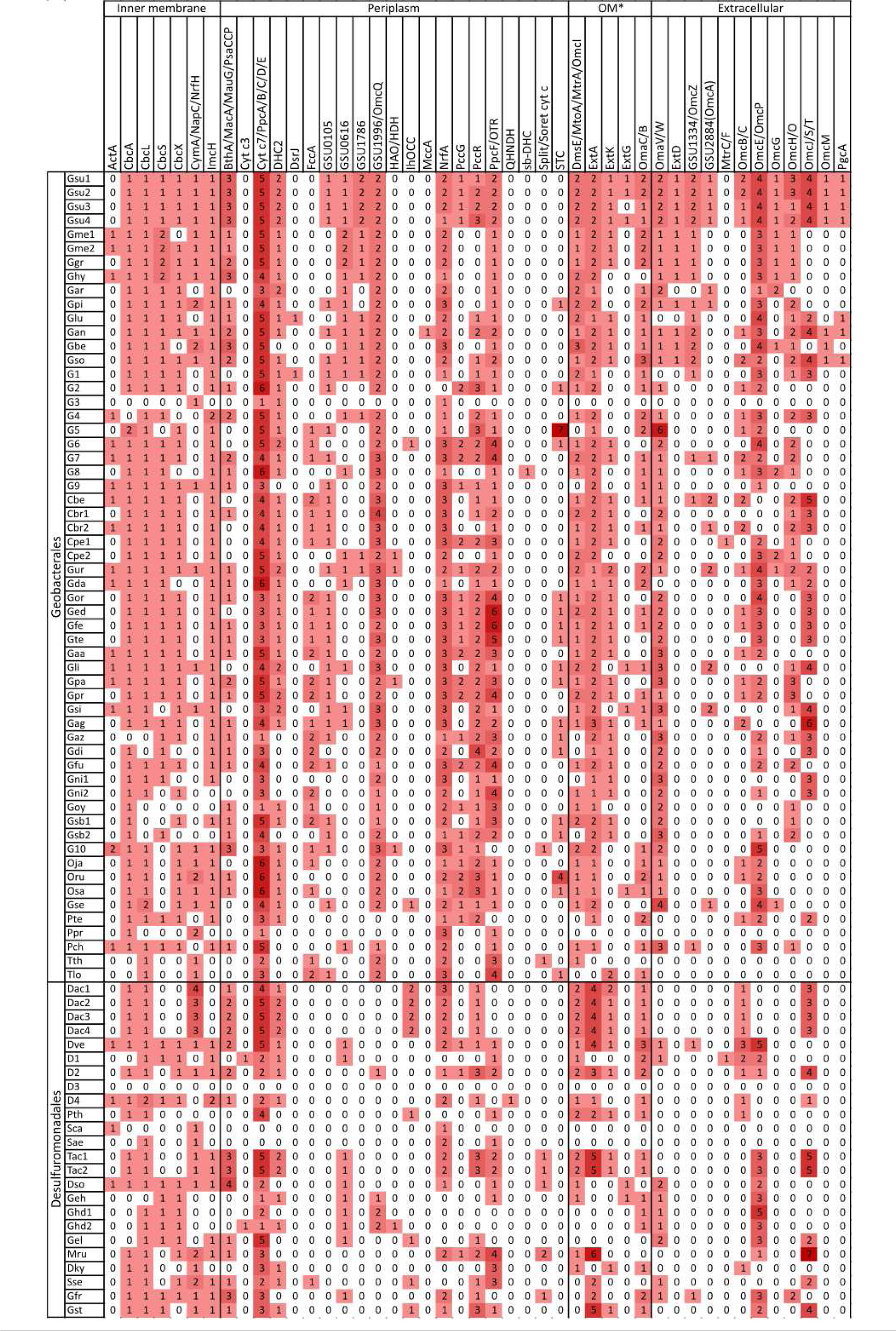
Heat map distribution of the different MHC across the species of the *Desulfuromonadia* class. The number of sequences found per homologous group is shown. High values are shown with darker red colors as the background. Species names are abbreviated and are shown in the same order as in Fig. 1 and 3. Raw data is provided in Data S4. Only DmsE/MtoA/MtrA/OmcI, ExtA, ExtK, ExtG, OmaC/B, OmaV/W (ExtC), ExtD, GSU2884(OmcA), MtrC/F, OmcB/C MHC are in operons coding for cytochrome-porin complexes. * Outer membrane.

Globally, *Desulfuromonadia* organisms exhibit diverse MHC sequences, featuring varying degrees of redundancy. Within these organisms, multiple parallel pathways seem to exist, at least as options at the level of the genome. This matches the observation in *G. sulfurreducens* and *S. oneidensis* of multiple pathways that offer alternative routes for reduction or oxidation of insoluble acceptors and donors, respectively (*66*, *78*), which is often linked to their ability in fine-tuning access to various redox potential ranges and conserve energy (*79*, *80*). Ultimately, this underscores the importance of having diverse MHC as a beneficial option to adapt to multiple conditions, as previously proposed (*81*).

### The unidentified MHC show that structure is more conserved than sequence

Since almost half of the MHC sequences remained unidentified due to a lack of characterized homologues, we explored the structural characteristics of the most representative groups. We assembled clusters based on sequence alignments and predicted the structure of representatives from clusters containing more than 15 sequences (41 clusters in total) using Alphafold2 (*82*). These conserved sequences are more likely to represent important proteins within *Desulfuromonadia* class, bridging the gaps in our current knowledge of their function and structural properties, and suggesting targets to be prioritized in future research. The most populated cluster, cluster 1, consists of 116 homologous sequences and the corresponding MHC are predicted to be located in the periplasmic space. The predicted structure reported in (Fig. 6A) is from a sequence from *Citrifermentans bemidjiense* Bem with eight heme-binding motifs. It distinctly displays the three long C-terminal alpha-helices and topological organization of the hemes typical of the group of octaheme cytochromes that catalyze key reactions in the nitrogen and sulfur cycles (*9*). Regarding the homologous octaheme MHC, the relative position of the distal ligands within this structure mirrors that of ONR, HAO/HDH and IhOCC (*9*), emphasizing that structure is more conserved than sequence. The second largest cluster displays a predicted structure that forms a long wire with closely spaced hemes similar to the already characterized nanowires (*11–14*, *36*) but forming a longer wire per monomer/polypeptide chain. This predicted structure has the closest neighboring hemes with predicted iron-to-iron distances lower than 15 Ångstroms, which is compatible with fast electron transfer along the length of the protein (*83*). As a representative, we have the predicted structure of a putative MHC from *Geobacter metallireducens* RCH3, featuring 27 heme-binding motifs and a molecular weight of 130.7 kDa (Fig 6B). The third largest cluster is predicted to be periplasmic, has 11 heme-binding motifs and the representative depicted in Fig. 6C is from *Desulfuromusa kysingii* with a predicted molecular weight of 51.3 kDa. Cluster 2 (Fig. 6B) is predicted to be extracellular and is not part of an operon coding for a porin-cytochrome complex. Proteins from this cluster have a predicted length that is more than the average width of the periplasm (350 Ångstroms vs 170 Ångstroms) (*84*). Maturation and translocation to the outer membrane of proteins of this size may pose potential challenges in the crowded periplasmic space. However, judging by the high prevalence of regions without secondary structural elements across the length of the protein we can expect a high degree of flexibility, potentially easing its processing into the mature form and delivery to the target cell location. Indeed, an architecture of modules connected by linkers that provide some flexibility may have been an evolved strategy to solve two problems. On the one hand, it limits the search space for distal ligands of the hemes to residues in each module, decreasing the risk of misfolding. On the other hand, the flexibility allows for easier navigation in the crowded periplasmic space when the mature protein needs to reach its final destination outside of the cell surface. Experimental characterization for a MHC with these characteristics was only achieved for GSU1996 that shows a wire-like structure, proposed to arise from the duplication and concatenation of cytochrome *c*_7_ modules (*36*, *85*). The molecular wire clusters identified here might have originated through similar processes, or GSU1996 could be the result of the pruning of larger ancestral cytochromes as found for other MHC (*9*). We provide an atlas of the remaining predicted structures from the representatives of the top 41 clusters in Fig. S2 (corresponding protein structure files in Data S5 and confidence estimation in Fig. S3), showing that there are likely still novel MHC folds that have not been characterized to date, including long tubular MHC, and MHC with novel beta-sheet domains. The NrfA heme arrangement seems to be highly common across different clusters (cluster 1, **<u>3</u>** and **<u>8</u>** – periplasm; clusters 4, 11 and 13 – unknown localization; Fig. S2). Some of these have 5 hemes and contain a C-terminal domain of beta sheets (cluster 4, 11 and 13 –unknown localization; Fig. S2) instead of alpha helixes as in NrfA (*5*). Fig. S2 also shows the presence of an extensive pool of periplasmic wire MHCs that begs the question of what is their physiological function, particularly considering that they have an average length similar to that generally associated with the width of the periplasm of Gram-negative bacteria (*86*). Either the periplasm of the corresponding *Desulfuromonadia* members is wider than average, or respiratory pathways in organisms that contain these putative MHC may possess features in the periplasm that are currently unknown. This MHC might directly link the two membranes, offering an alternative to small soluble cytochromes that depend on diffusion for encountering partner proteins to engage in electron transfer from the quinone pool to the cell surface. The top three MHC clusters with a representative predicted to have an extracellar location showed the presence of one or more hematite-binding motifs (*87*) that were surface exposed and near a heme. From these, cluster 1 was predicted to form a long nanwire wih 27 hemes and displays this feature unlike what is observed for the structurarly characterized extracellular nanowires OmcS, OmcE and OmcZ of *Geobacter sulfurreducens* (*11–13*). Among the MHC predicted to have an extracellular location and that were characterized only on the basis of genetics, GSU2884 (OmcA), OmcH, OmcG, OmcM also showed the presence of one or multiple hematite-binding motifs. From these, GSU2884 (OmcA), OmcH, OmcG are also predicted to form long naowires with high number of hemes per polipetide chain (Fig. S1). Fig. 5 shows that species of the *Desulfuromonadia* class contain MHC that are typically associated with *Shewanella* spp. and other related Gamma-proteobacteria, namely FccA, STC, DmsE/MtoA/MtrA/OmcI MtrC/F. From the pool of unidentified sequences based on sequence alignments, we also found structures resembling MtrC/F/OmcA like MHC (Fig. S2; the second largest cluster of extracellular MHC composed of 57 sequences), which are not associated with cytochrome-porin complexes according to the operon configuration from the reference sequence. Indeed, in *Shewanella oneidensis* MR-1, OmcA and UndA that also belong to this family are also not part of the same operon (*88*, *89*). Further investigation is however required to understand if these sequences derive from vertical or horizontal gene transfer. Several studies point out that MHC have a high likelihood of horizontal gene transfer, including some of those extensively characterized in *Shewanella* spp. (*90*). Alternatively, these could be distantly related MHC derived from a common ancestor of the two phyla to which the *Desulfuromonadia* and *Shewanella* belong.

**Fig 6.**
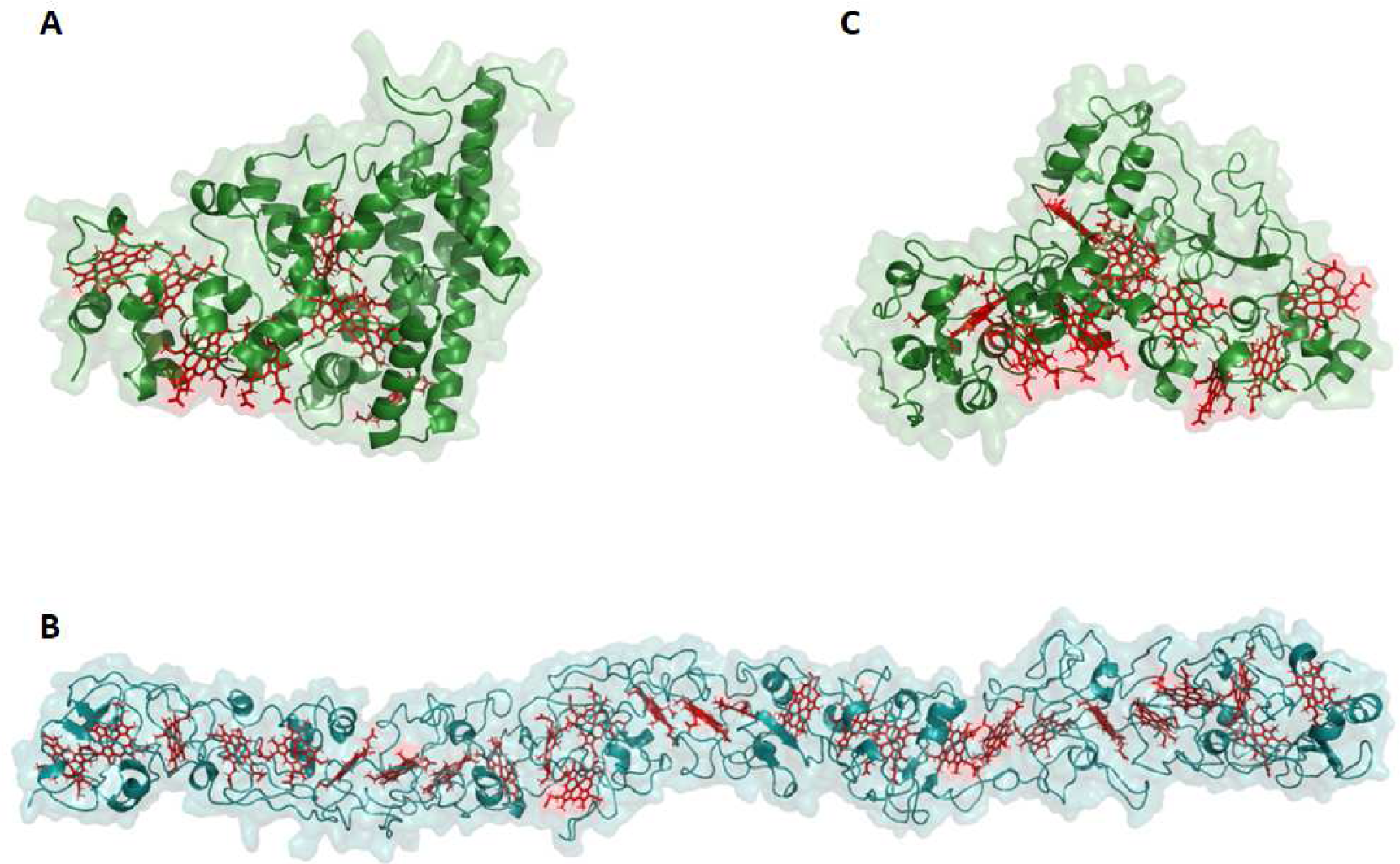
Predicted structures from the top three representative clusters for which no experimentally determined structure is available. Structures are oriented with the N-terminal at the left-hand side. (A) Structure representative of the largest cluster whose sequence is from *Citrifermentans bemidjiense* and is predicted to be located in the periplasm. (B) Structure representative of the second largest cluster whose sequence is from *Geobacter metallireducens* and is predicted to be located at the extracellular space. (C) Structure representative of the third largest cluster whose sequence is from *Desulfuromusa kysingii* and is predicted to be located in the periplasm.

### MHC diversity correlates with subcellular location

From the 4716 MHC sequences of the dataset, 51% could be assigned to a specific cellular sublocation. From these 11.1% were assigned to the inner membrane, 62.8 % to the periplasm, 0.1% to the outer membrane and 25.9% to the extracellular space. Fig. 7 shows that inner membrane MHC are more conserved with respect to size and number of hemes, while extracellular MHC are the least conserved. The inner membrane MHC group only contains three sequences that have more than 10 heme-binding motifs. Conversely, the extracellular group of sequences contains elements ranging from 2 to 75 heme-binding motifs, which are well distributed at much higher numbers of heme-binding motifs. In this group, 50% of the sequences have 10 or more hemes. MHC sequences assigned to the periplasm are also well-distributed towards relatively high numbers of heme-binding motifs. Only a few sequences were assigned to the outer membrane region. This can be a consequence of the fact that the distinction between outer membrane and periplasmic location can be difficult to establish. For example, some outer membrane MHC such as MtrA are also proposed to function in the periplasm (*91*, *92*). The pattern of diversification, in terms of size and number of heme binding sites versus cell location, shows that there is more room for diversification at the cell surface than on the inner membrane. This likely reflects the fact that extracellular MHC confer an adaptative advantage to these organisms in diverse ecological contexts, whereas MHC of the inner-membrane are functionally more homogeneous and involved in metabolic functions that are conserved across the organisms of the class, namely exchanging electrons between the quinone/quinol pool and the periplasmic redox partners, and translocation of protons across the inner-membrane for energy conservation (*46*, *50*). In this sense, evolution appears to have more leeway to modify extracellular MHC, accommodating multiple paralogues and changing the sequence and structure of these MHC, leading to a high degree of diversity.

**Fig. 7.**
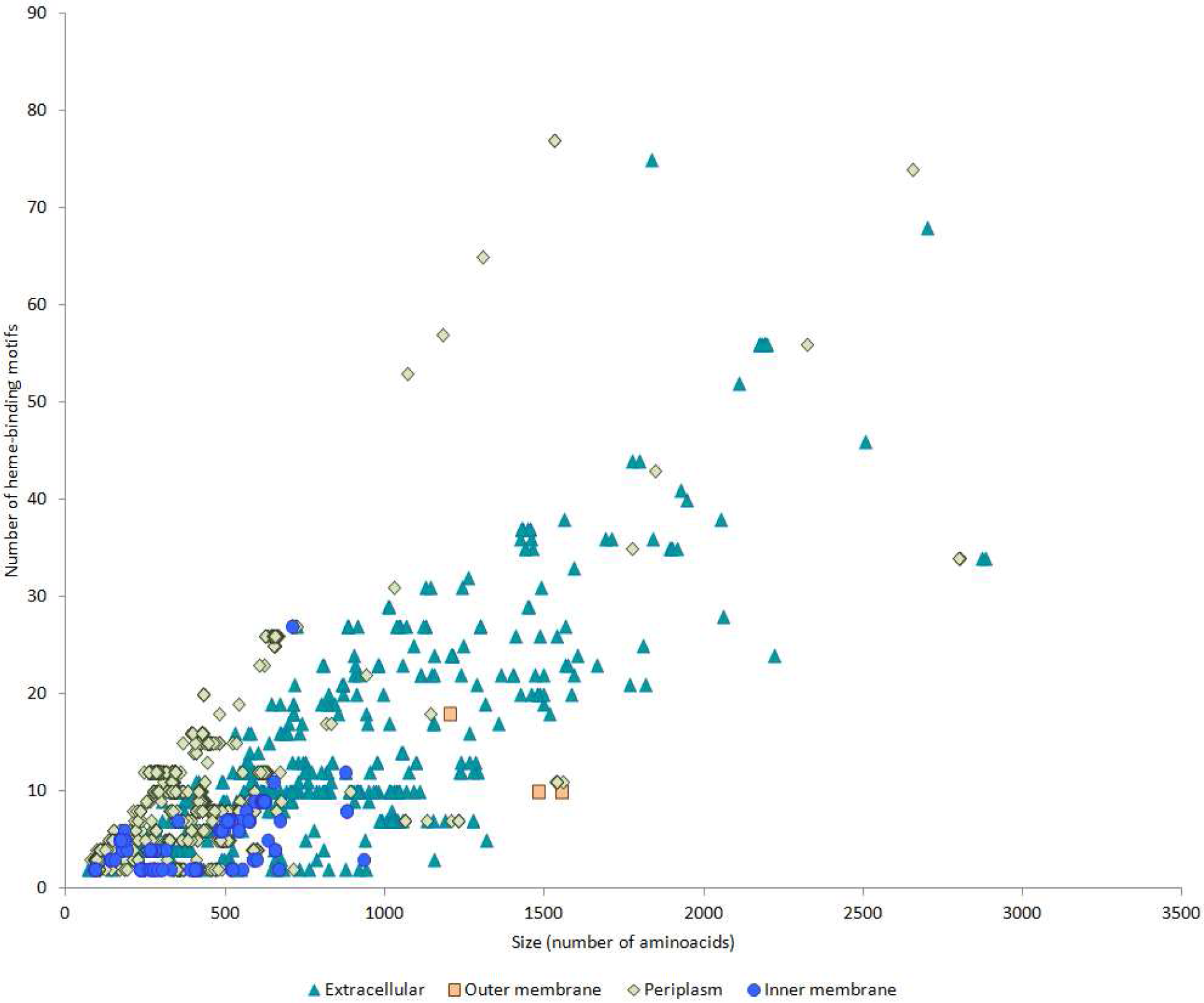
Plot of size vs number of hemes in the MHC sequences for which a subcellular location could be assigned.

## Discussion

MHC are key metalloproteins that intervene in the biogeochemical cycles of the elements. Even though MHC have been characterized in great detail for almost half a century, experimental characterization has only focused on a few representatives from the vast universe of MHC. Given that *Desulfuronadia* class have an average prevalence of putative MHC in their genomes that is larger than what is typical for other prokaryotes, it was targeted to expand our understanding of this important class of proteins. This study revealed for the first time the diversity of MHC of the *Desulfuromonadia* class that contains metabolically diverse organisms with fermentative and/or respiratory metabolisms. Those that have a respiratory energetic metabolism have a relatively high number of MHC per genome, with the highest numbers found in some members of the *Geobacterales* order. The number of heme-binding motifs per polypeptide chain is highly variable reaching numbers that are several fold higher than the existing maximum of the MHC currently characterized structurally. We presently have limited knowledge about the underlying reasons for the distinct option between oligomerization and repetition of polypeptide chain modules in modular evolution. Nevertheless, the frequency of heme-binding motifs is concentrated towards relatively low numbers. This shows biological constraints that only allow the number of heme-binding motifs to reach a relatively high number on very specific occasions. Experimental work is still far from giving answers on what factors come to bear on the solutions found by nature on these specific occasions. Nevertheless, this study reinforces the need to focus on this matter in order to be able to rationally manipulate MHC without being limited to using MHC with a relatively low number of hemes for biotechnological applications, given that it has been known for almost half a century that these proteins have interesting electrical properties (*93*). MHC from the different cellular locations show different patterns of conservation. Inner membrane MHC even though having low relative abundance in total, present higher conservation. By contrast, extracellular MHC are the least conserved in general. These observations correlate with the particular environment of the two cellular locations. Whereas innermembrane MHC for the most part, interact with a homogeneous set of conditions involving electron exchange with the quinone pool, outer-membrane and extracellular MHC evolved to match the diversity of environmental conditions of the ecosystem of the organism to which they belong. However, even for inner-membrane MHC, we still lack detailed characterization of some of these for the model organisms that perform extracellular electron transfer. Furthermore, this work reveals that we lack diversity of characterized MHC in general, where it was only possible to identify half of the MHC sequences from the *Desulfuromonadia*. This observation reinforces the potential of the so-called protein structuromics in leveraging structural prediction tools to explore the role of proteins that have not been characterized biochemically yet, and identify interesting targets for this characterization (*94*). On the one hand, some uncharacterized examples represent novel nanowire-like MHC that have higher molecular weight and heme content than the currently characterized nanowire MHC. On the other hand, some of the sequences that did not fall into recognized MHC families turned out to have predicted structures that match known families. This shows that just as there is redundancy in the genetic code when translated to protein sequence, there is redundancy in the sequence code when translated to structure. This redundancy also translates to functional redundancy as is known for some MHC families such as that containing NrfA, where some members have diverse potential catalytic activities (*9*). This provides a mechanism for the safe transition between different functions as natural selection operates on existing proteins to provide additional fitness to the hosting organism in a changing environment. Overall, this study reveals the diversity of MHC of *Desufuromonadia* with respect to their structure, function, and evolutionary plasticity to match diverse environmental conditions. It shows that evolution has found more diverse solutions for extracellular exposed MHC, in agreement with the need to match diverse environmental conditions that may in some cases approach the minimum energetic needs to sustain the metabolism (*81*). This information broadens our capability to understand the functional role of individual MHC and to improve their properties to match artificial environments such as those found in microbial electrochemical technologies.

## Materials and Methods

### Data acquisition and distribution of MHC and heme-binding motifs across the Desulfuromonadia class

Genomes of *Desulfuromonadia* were obtained from the RefSeq database (*95*) using the NCBI genome server on 5^th^ of April, 2022. All genomes assigned to ‘*Desulfuromonadales’* were downloaded given that NCBI still had the previous classification that is now further substituted by the class *Desulfuromonadia*. When genomes from the same strain existed, the one which had a higher level of assembly was chosen. If the same level of assembly was present, the most recent genome submission was chosen. A list of the genomes and their global characteristics is shown in Data S1. Corresponding protein-coding sequences were obtained from each genome and text manipulation tools of Ubuntu 18.04 (Bash version 4.4.19(1)) were used to tag each protein sequence to the corresponding genome reference number. MHC were identified using InterproScan 5.55-88.0 (*96*). NCBI eliminates redundancy from dataset copies where paralogues with 100% identity are assigned to only one reference code and proteome datasets only contain one entry in this case. To circumvent this and take into account MHC identical copies in the genome, each genome was searched for copies using Ubuntu 18.04 (Bash version 4.4.19(1)). When there were copies of MHC these were further added to the MHC protein coding sequences dataset. MHC sequences were converted to table format and transferred to Microsoft Excel, where heme-binding motifs were counted using the wildcard pattern recognition function. The canonical CXXCH and alternative heme-binding motifs were searched (*4*, *9*). Sequences showing less than two putative heme-binding motifs were removed. Graphs for the number of MHC per gnome and the global distribution of the number of heme-binding motifs across all genomes were generated using Microsoft Excel. Box-plots corresponding to the genomes’ individual distributions of the number of heme-binding motifs were generated using BoxPlotR (*97*). MHC localization was predicted using PSORTb 3.0.3 (*98*). Molecular weight was predicted using Expasy PI/MW tool (*99*) and adding 616.18 Da for each heme.

### Identification, clustering and structure prediction of MHC

A curated database of characterized MHC was constructed using the table reported in the literature (*2*) with further additions to update the list of MHC. The source databases from which these sequences were collected were the NCBI-RefSeq (*95*) and PDB. Each sequence was aligned against each sequence of this database using BLASTp suite-2sequences (Tatusova and Madden, 1999) with a cutoff of 1^-5^ *e*-value. Alignment data were retrieved and query and subject length and coverage were calculated using Microsoft Excel. Alignments were then further filtered using a minimum of 70% query and subject coverage, 30% length difference between query and subject and a minimum of 25% identity. Sequences from the constructed database were clustered using MMseqs 2 (*101*) with 25% identity, 70% coverage with coverage mode 0 (which applies this threshold to both query to subject and subject to query) in order to group them as homologous groups. Each sequence that matched the alignment criteria was labeled with the subject protein sequence from the constructed database. In addition, each sequence was also labeled with the homologous group where that subject sequence belonged. Unidentified MHC were further clustered using MMseqs 2 (*101*) with 20% identity, 70% coverage with coverage mode 0. Predicted structures were generated using AlphaFold2 (2.3.2)(*82*) within the COSMIC2 platform (*102*), using as parameters the full database sequence for the multiple sequence alignments (MSA), five predictions per model, monomer mode for the Predicted Template Modeling Score (pTM) and Amber relaxation for the best model generated. For the characterized homologous groups from which no representative structure is available in the PDB, the predicted structure was gathered from the AlphaFold Protein Structure database (*103*). For structure prediction the signal peptide was cut using SignalIP 5.0 (*104*). To incorporate hemes into the structural models, a PyMOL script was devised. In brief, it operates by identifying CXXCH and CXXXXCH binding motifs and then iterates the PyMOL align function to incorporate heme 3 from PDB entry 6HR0 at those binding motifs. This script has now been made available at https://github.com/BenjaminNash5/C_heme_insert_script_repository. The structures with the hemes were subsequently water-refined using the protocol implemented in the refinement interface of the HADDOCK 2.4 webserver (*105*, *106*). Image processing was performed using PyMOL Molecular Graphics System (version 2.0 Schrödinger, LLC).

## Acknowledgments

The authors are grateful to Tom Clarke for helpful suggestions

## Funding

Financial support was provided by the following institutes and funding agencies with the respective projects specified

MOSTMICRO-ITQB base funding with references UIDB/04612/2020 and UIDP/04612/2020

LS4FUTURE Associated Laboratory (LA/P/0087/2020)

Portuguese Foundation for Science and Technology (FCT) grant PTDC/BIABQM/4143/2021.

## Author contributions

Conceptualization: RS, BMF, CMP, ROL

Methodology: RS, BMF, BWN, ROL

Investigation: RS, BMF, ROL

Visualization: RS

Supervision: CMP, ROL

Writing original draft: RS

Writing review & editing: RS, BMF, BWN, CMP, ROL

## Data and materials availability

All relevant data are available in the supplementary materials. The PyMoL script for heme *c* incorporation within protein structure files is available through this repository: https://github.com/BenjaminNash5/C_heme_insert_script_repository

## Supplementary Materials

