## Supplementary material for "A Survey of the *Desulfuromonadia* cytochromome provides a glimpse of the unexplored diversity of multiheme cytochromes in nature": suplementary material

Ricardo Soares *et al.*

**This PDF file includes:**

Figs. S1 to S3
Tables S1
Data S1 to S5

**Inner membrane**

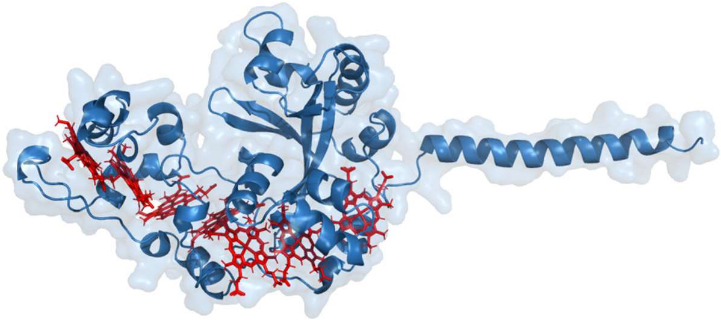

Protein: CbcA
Organism: *Geobacter sulfurreducens*
Number of heme-binding motifs: 7
Predicted molecular weight: 40.2 kDa

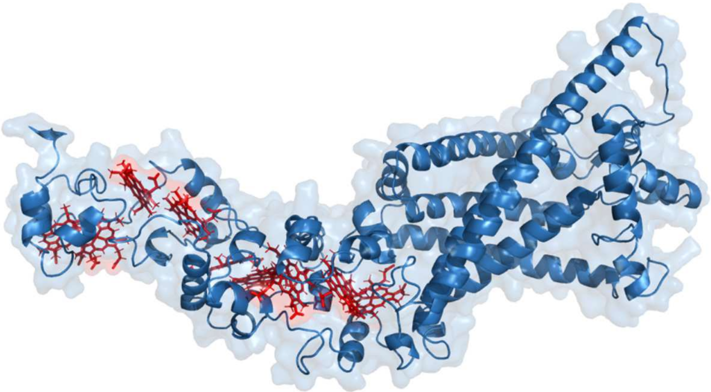

Protein: CbcL
Organism: *Geobacter sulfurreducens*
Number of heme-binding motifs: 9
Predicted molecular weight: 73.3 kDa

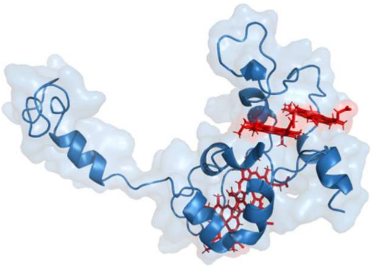

Protein: CbcS
Organism: *Geobacter sulfurreducens*

Number of heme-binding motifs: 4
Predicted molecular weight: 17.9 kDa

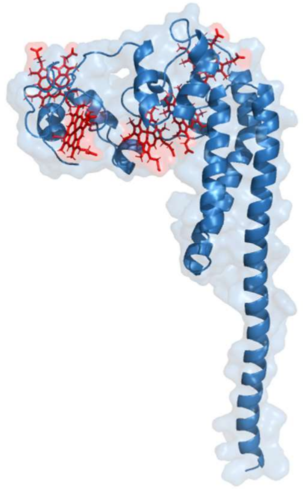

Protein: CbcX
Organism: *Geobacter sulfurreducens*
Number of heme-binding motifs: 5
Predicted molecular weight: 28.8 kDa

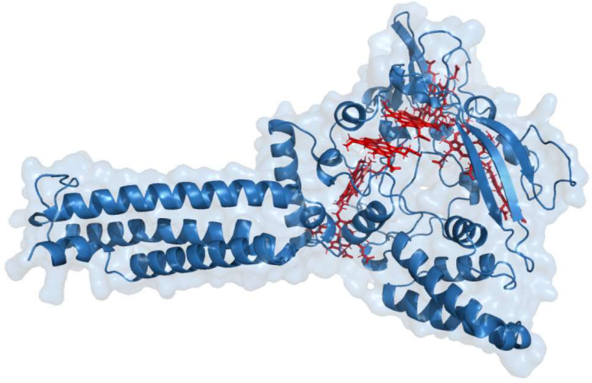

Protein: ImcH
Organism: *Geobacter sulfurreducens*
Number of heme-binding motifs: 7
Predicted molecular weight: 61.7 kDa

**Periplasm**

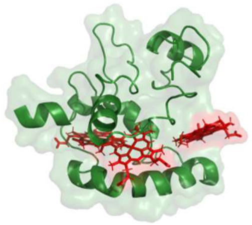

Protein: DsrJ
Organism: *Allochromatium vinosum* ATCC 17899
Number of heme-binding motifs: 3
Predicted molecular weight:  15.5 kDa

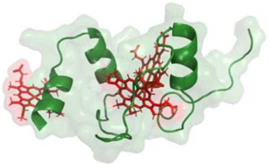

Protein: GSU0105
Organism: *Geobacter sulfurreducens*
Number of heme-binding motifs: 3
Predicted molecular weight: 9.6 kDa

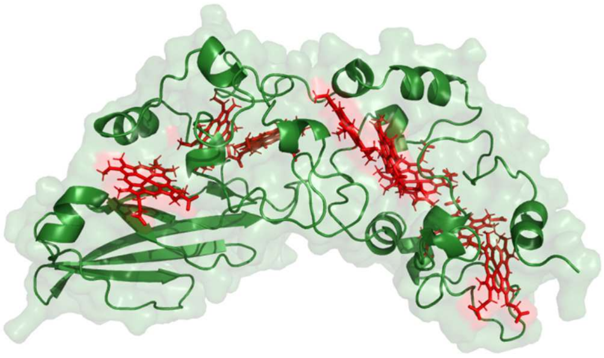

Protein: GSU0616
Organism: *Geobacter sulfurreducens*
Number of heme-binding motifs: 8
Predicted molecular weight: 38.8 kDa

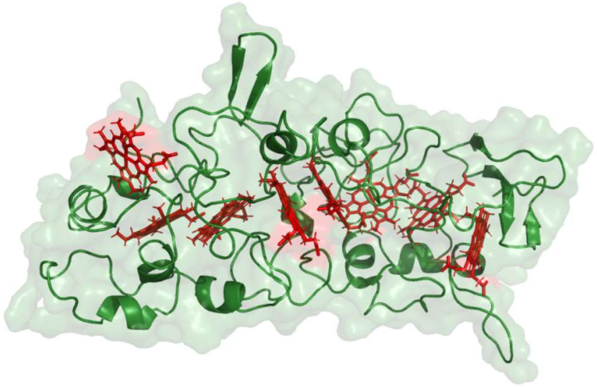

Protein: GSU1786
Organism: *Geobacter sulfurreducens*
Number of heme-binding motifs: 8
Predicted molecular weight: 42.8 kDa

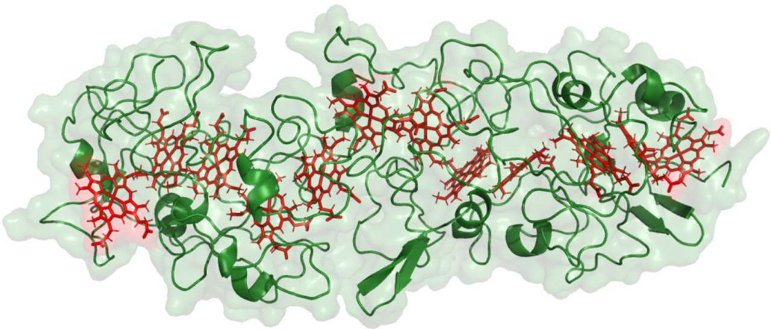

Protein: PccG
Organism: *Geobacter sulfurreducens*
Number of heme-binding motifs: 12
Predicted molecular weight: 66.9 kDa

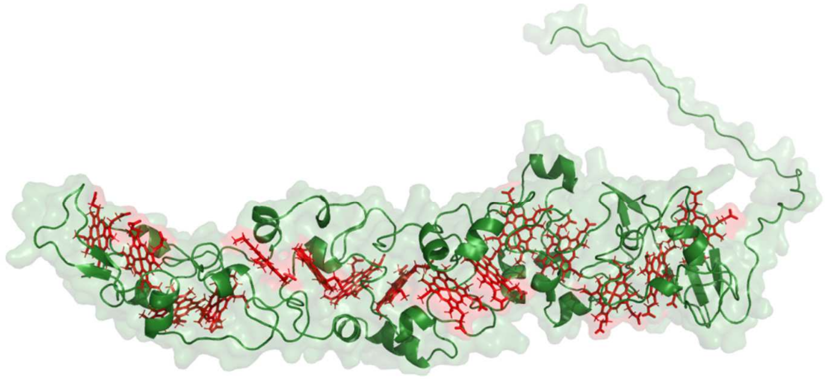

Protein: PccR
Organism: *Geobacter sulfurreducens*
Number of heme-binding motifs: 15
Predicted molecular weight: 59.4 kDa

**Outer membrane**

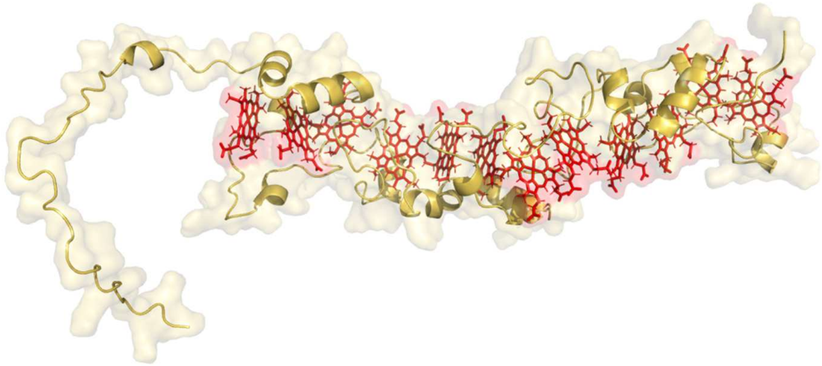

Protein: ExtA
Organism: *Geobacter sulfurreducens*
Number of heme-binding motifs: 12
Predicted molecular weight: 37.8 kDa

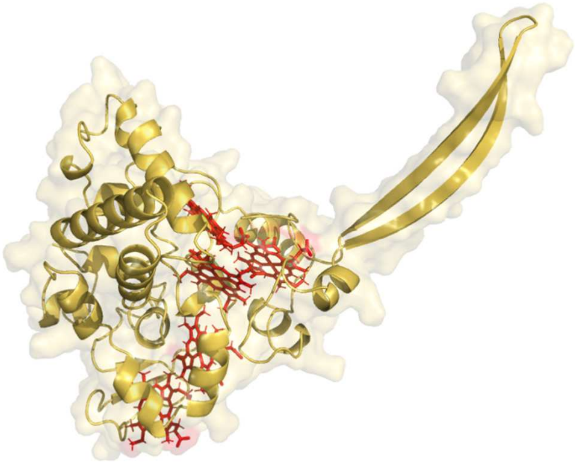

Protein: ExtK
Organism: *Geobacter sulfurreducens*
Number of heme-binding motifs: 5
Predicted molecular weight: 35.8 kDa

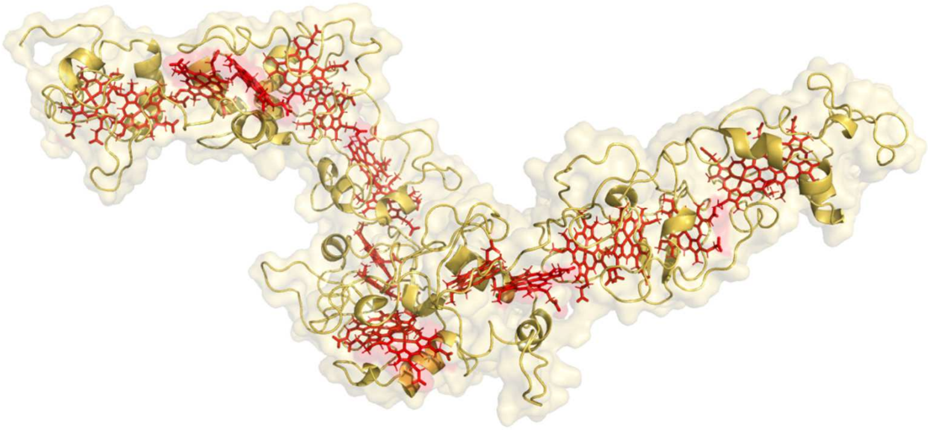

Protein: ExtG
Organism: *Geobacter sulfurreducens*
Number of heme-binding motifs: 18
Predicted molecular weight: 79.5 kDa

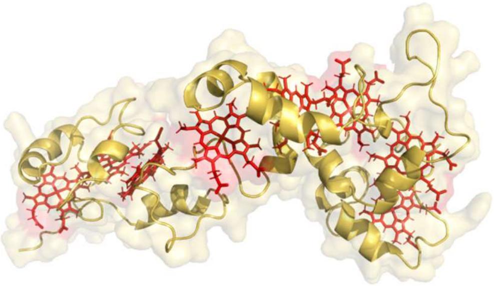

Protein: OmaB
Organism: *Geobacter sulfurreducens*
Number of heme-binding motifs: 8
Predicted molecular weight: 28.1 kDa

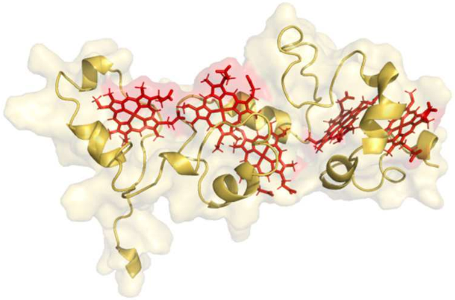

Protein:  OmaW
Organism: *Geobacter sulfurreducens*

Number of heme-binding motifs: 5
Predicted molecular weight: 18.9 kDa

**Extracellular**

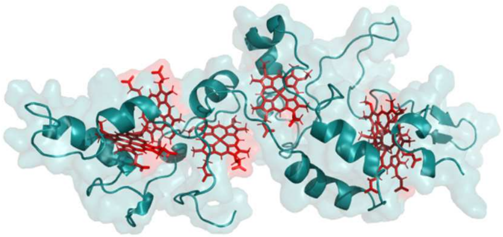

Protein: ExtD
Organism: *Geobacter sulfurreducens*
Number of heme-binding motifs: 6
Predicted molecular weight: 26.8 kDa

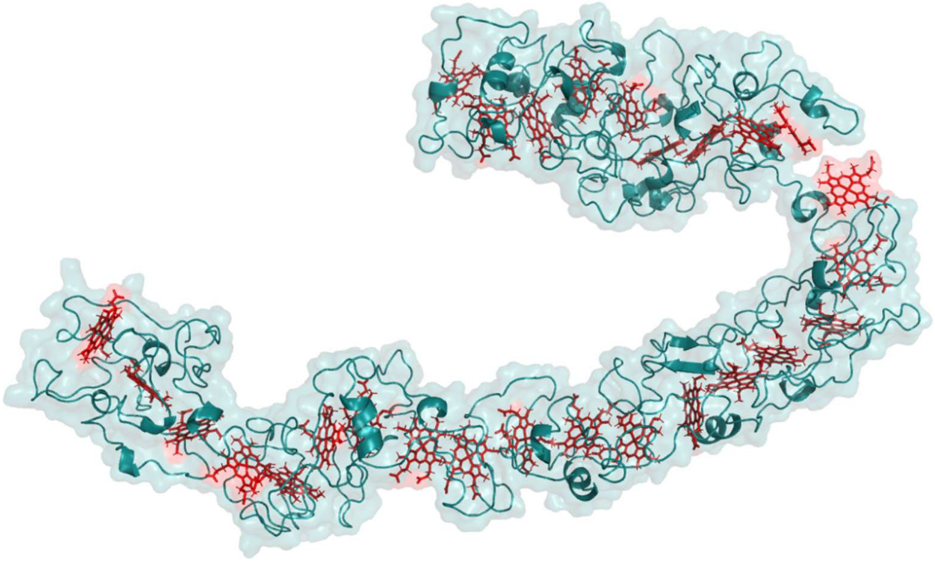

Protein: GSU2884 (OmcA)
Organism: *Geobacter sulfurreducens*
Number of heme-binding motifs: 27
Predicted molecular weight: 117.9 kDa

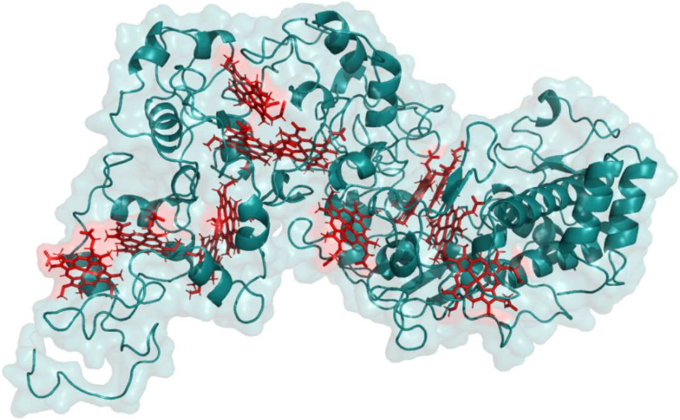

Protein: OmcB
Organism: *Geobacter sulfurreducens*
Number of heme-binding motifs: 12
Predicted molecular weight: 82.2 kDa

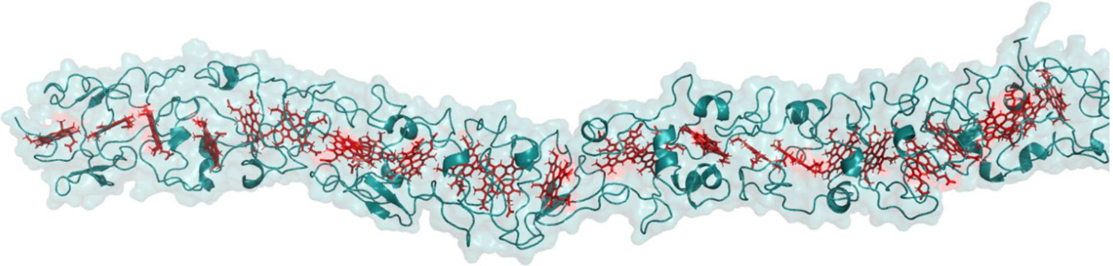

Protein: OmcH
Organism: *Geobacter sulfurreducens*
Number of heme-binding motifs: 24
Predicted molecular weight:  102.9 kDa

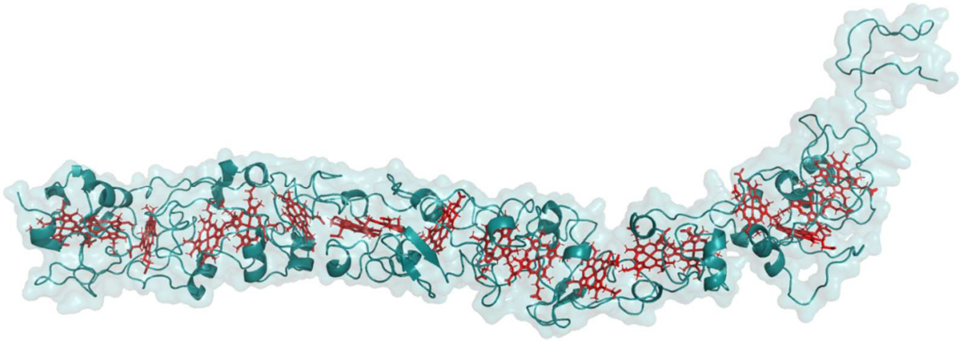

Protein: OmcG
Organism: *Geobacter sulfurreducens*
Number of heme-binding motifs: 19
Predicted molecular weight: 79.2 kDa

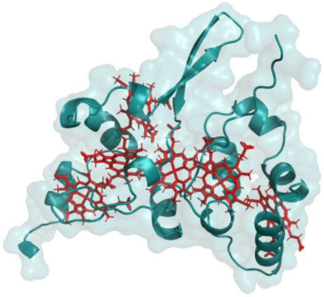

Protein: OmcM
Organism: *Geobacter sulfurreducens*
Number of heme-binding motifs: 6
Predicted molecular weight: 19.9 kDa

**Fig. S1.**

AlphaFold2 structure prediction of the MHC used as characterized references from which no available structure is present. Organism from which each sequence belongs, number of heme-binding motifs and the predicted molecular weight (hemes included) is shown bellow each predicted structure.

**Inner membrane**

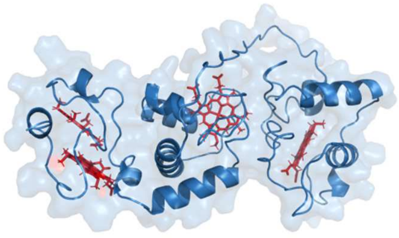

**Cluster: 1**
Sequence: WP_149309508.1
Cluster size: 49
Organism: *Oryzomonas rubra*
Number of heme-binding motifs: 4
Predicted molecular weight: 29.0 kDa

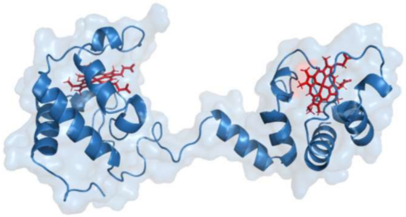

**Cluster: 2**
Sequence: WP_039642720.1
Cluster size: 29
Organism: *Geobacter anodireducens*
Number of heme-binding motifs: 2
Predicted molecular weight: 23.2 kDa

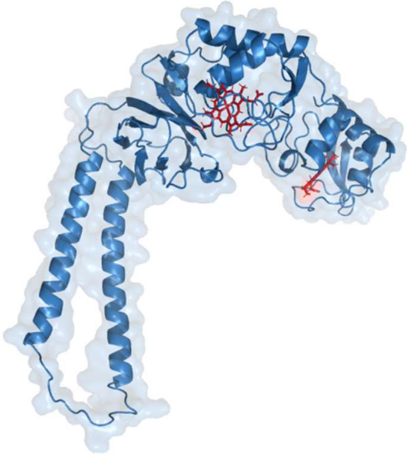

**Cluster: 3**
Sequence: WP_221249441.1
Cluster size: 19
Organism: *Desulfuromonas versatilis*
Number of heme-binding motifs: 2
Predicted molecular weight: 46.1 kDa

**Periplasm**

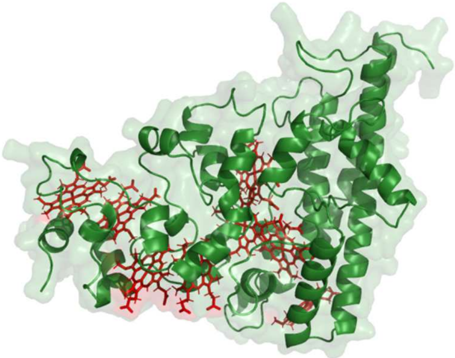

**Cluster: 1**
Sequence: WP_012532096.1
Cluster size: 116
Organism: *Citrifermentans bemidjiense*
Number of heme-binding motifs: 8
Predicted molecular weight: 47.7 kDa

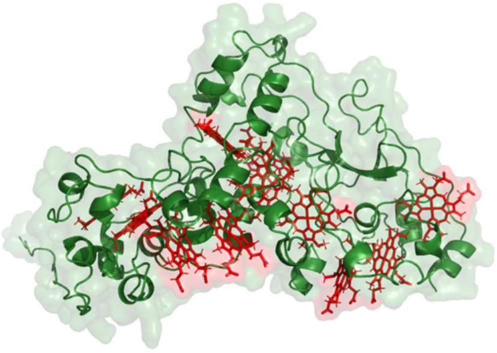

**Cluster: 2**
Sequence: WP_092345198.1
Cluster size: 69
Organism: *Desulfuromusa kysingii*
Number of heme-binding motifs: 11
Predicted molecular weight: 51.3 kDa

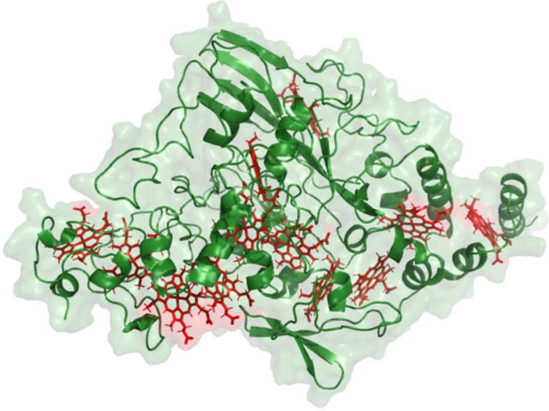

**Cluster: 3**
Sequence: WP_145019020.1
Cluster size: 53
Organism: *Geobacter argillaceus*
Number of heme-binding motifs: 12
Predicted molecular weight: 71.9 kDa

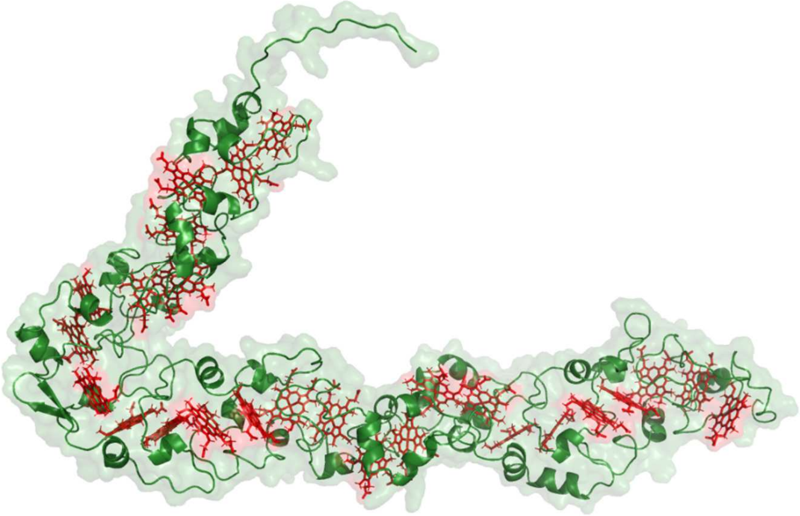

**Cluster: 4**
Sequence: WP_191156322.1
Cluster size: 49
Organism: *Pelovirga terrestris*
Number of heme-binding motifs: 26
Predicted molecular weight: 82.3 kDa

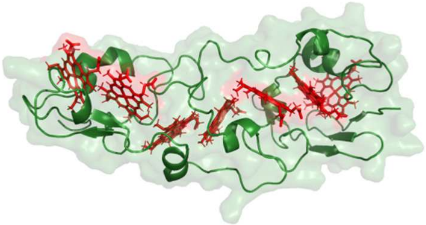

**Cluster: 5**
Sequence: WP_230997429.1
Cluster size: 33
Organism: *Geomonas* sp. RF6
Number of heme-binding motifs: 7
Predicted molecular weight: 25.8 kDa

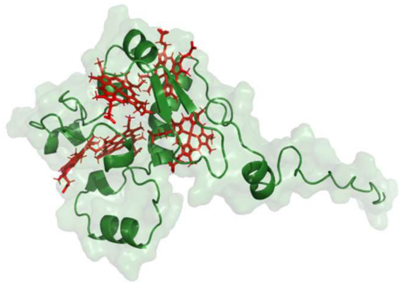

**Cluster: 6**
Sequence: WP_232278942.1
Cluster size: 28
Organism: *Geotalea uraniireducens*
Number of heme-binding motifs: 5
Predicted molecular weight: 19.2 kDa

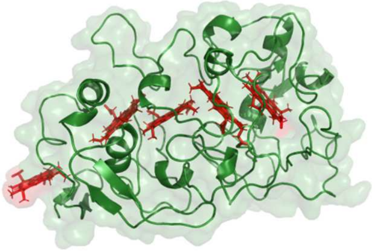

**Cluster: 7**
Sequence: WP_236685653.1
Cluster size: 20
Organism: *Geobacter pickeringii*
Number of heme-binding motifs: 5
Predicted molecular weight: 37.9 kDa

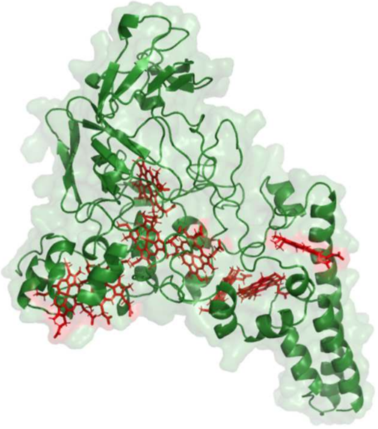

**Cluster: 8**
Sequence: WP_168205920.1
Cluster size: 18
Organism: *Geobacter* sp. FeAm09
Number of heme-binding motifs: 8
Predicted molecular weight: 58.1 kDa

**Extracellular**

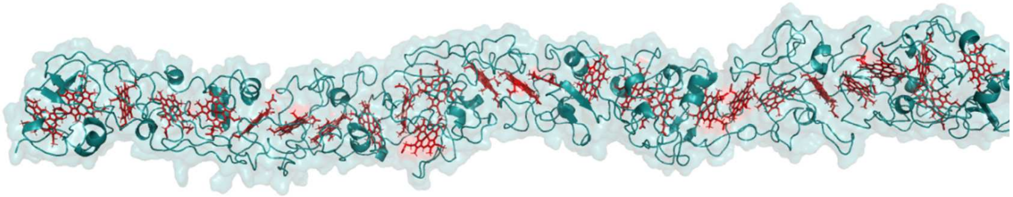

**Cluster: 1**
Sequence: WP_004512272.1
Cluster size: 104
Organism: *Geobacter metallireducens*
Number of heme-binding motifs: 27
Predicted molecular weight: 130.6 kDa

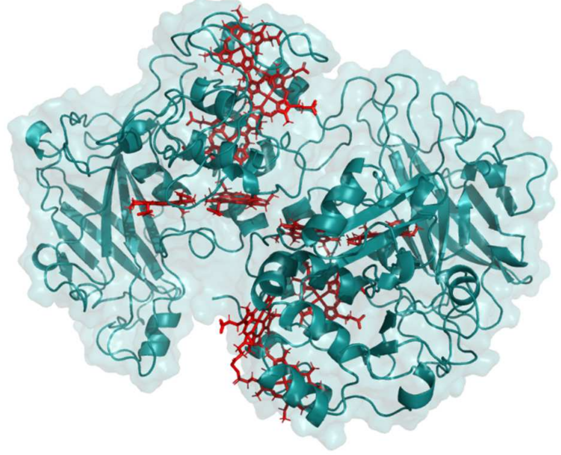

**Cluster: 2**
Sequence: WP_214172253.1
Cluster size: 57
Organism: *Citrifermentans pelophilum*
Number of heme-binding motifs: 10
Predicted molecular weight: 88.3 kDa

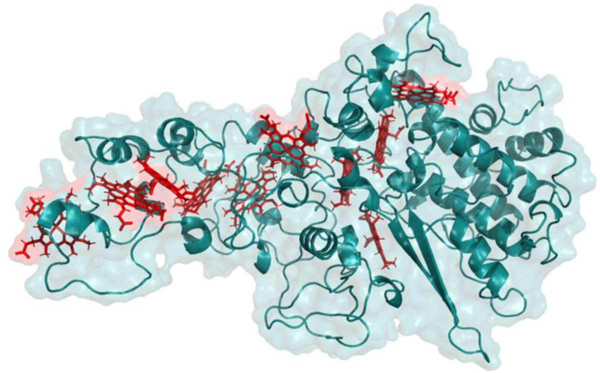

**Cluster: 3**
Sequence: WP_149211077.1
Cluster size: 56
Organism: *Geobacter* sp. FeAm09
Number of heme-binding motifs: 10
Predicted molecular weight: 70.0 kDa

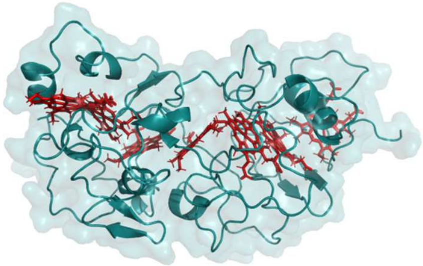

**Cluster: 4**
Sequence: WP_149208723.1
Cluster size: 33
Organism: *Geobacter* sp. FeAm09
Number of heme-binding motifs: 7
Predicted molecular weight: 40.0 kDa

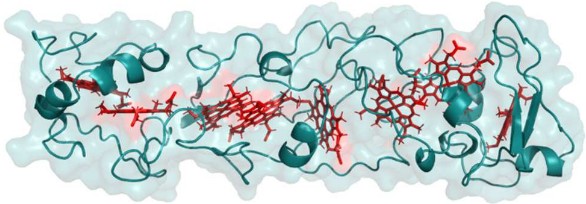

**Cluster: 5**
Sequence: WP_185244360.1
Cluster size: 16
Organism: *Citrifermentans bremense*
Number of heme-binding motifs: 8
Predicted molecular weight: 34.5 kDa

**Unknown localization**

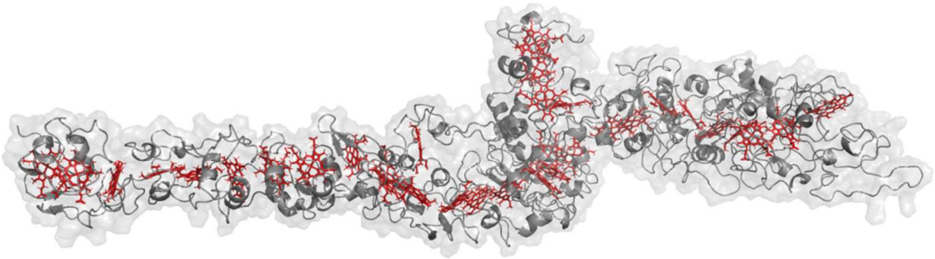

**Cluster: 1**
Sequence: WP_214184651.1
Cluster size: 66
Organism: *Geobacter hydrogenophilus*
Number of heme-binding motifs: 33
Predicted molecular weight: 178.9 kDa

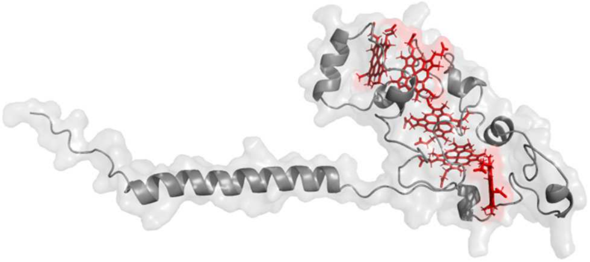

**Cluster: 2**
Sequence: WP_183348177.1
Cluster size: 46
Organism: *Geomonas paludis*
Number of heme-binding motifs: 5
Predicted molecular weight: 21.1 kDa

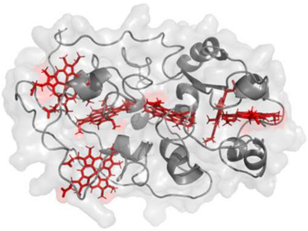

**Cluster: 3**
Sequence: WP_012645855.1
Cluster size: 45
Organism: *Geotalea daltonii*
Number of heme-binding motifs: 6
Predicted molecular weight: 29.6 kDa

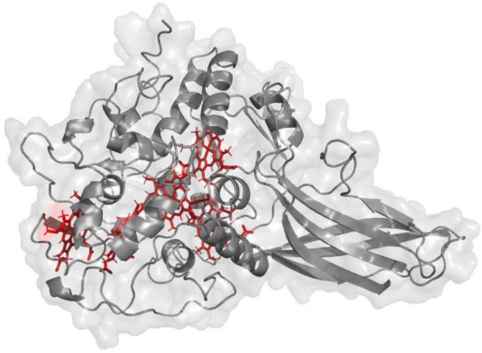

**Cluster: 4**
Sequence: WP_149210553.1
Cluster size: 40
Organism: *Geobacter* sp. FeAm09
Number of heme-binding motifs: 5
Predicted molecular weight: 53.3 kDa

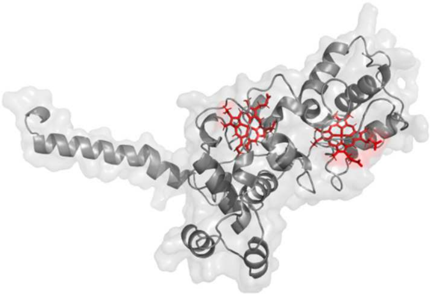

**Cluster: 5**
Sequence: WP_173199422.1
Cluster size: 38
Organism: *Geobacter* sp. SVR
Number of heme-binding motifs: 2
Predicted molecular weight: 33.4 kDa

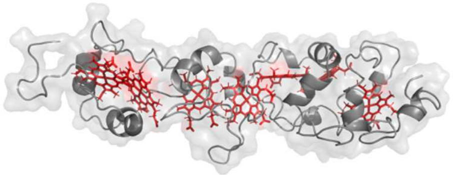

**Cluster: 6**
Sequence: WP_020675252.1
Cluster size: 38
Organism: *Geopsychrobacter electrodiphilus*
Number of heme-binding motifs: 7
Predicted molecular weight: 28.5 kDa

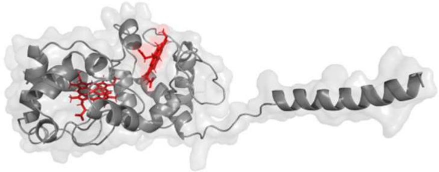

**Cluster: 7**
Sequence: WP_214186522.1
Cluster size: 36
Organism: *Geobacter hydrogenophilus*
Number of heme-binding motifs: 2
Predicted molecular weight: 26.4 kDa

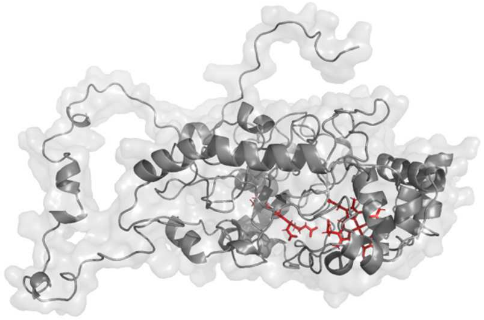

**Cluster: 8**
Sequence: WP_183354258.1
Cluster size: 34
Organism: *Geomonas silvestris*
Number of heme-binding motifs: 2
Predicted molecular weight: 57.4 kDa

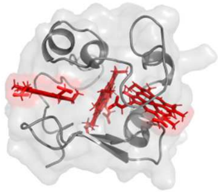

**Cluster: 9**
Sequence: WP_029912551.1
Cluster size: 33
Organism: *Pelobacter seleniigenes*
Number of heme-binding motifs: 3
Predicted molecular weight: 10.9 kDa

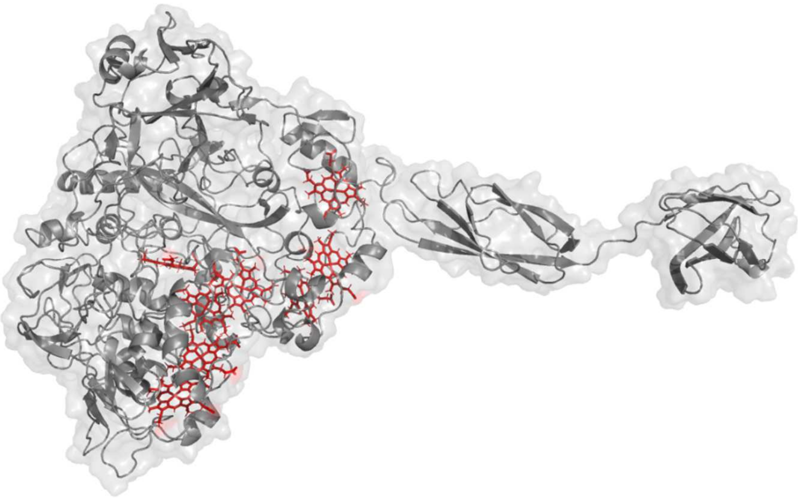

**Cluster: 10**
Sequence: WP_208610118.1

Cluster size: 32
Organism: *Malonomonas rubra*
Number of heme-binding motifs: 8
Predicted molecular weight: 123.6 kDa

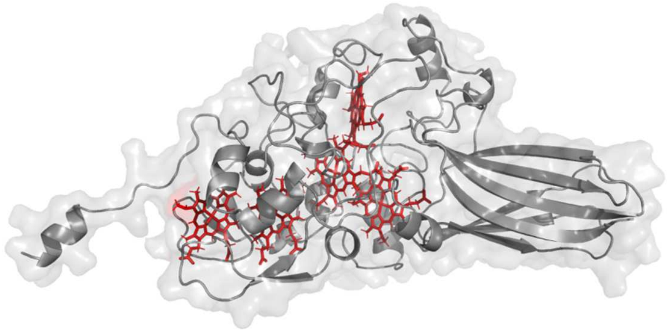

**Cluster: 11**
Sequence: WP_214296179.1
Cluster size: 32
Organism: *Geobacter chapellei*
Number of heme-binding motifs: 5
Predicted molecular weight: 46.4 kDa

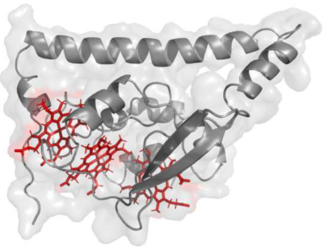

**Cluster: 12**
Sequence: WP_010943832.1
Cluster size: 28
Organism: *Geobacter sulfurreducens*
Number of heme-binding motifs: 3
Predicted molecular weight: 21.1 kDa

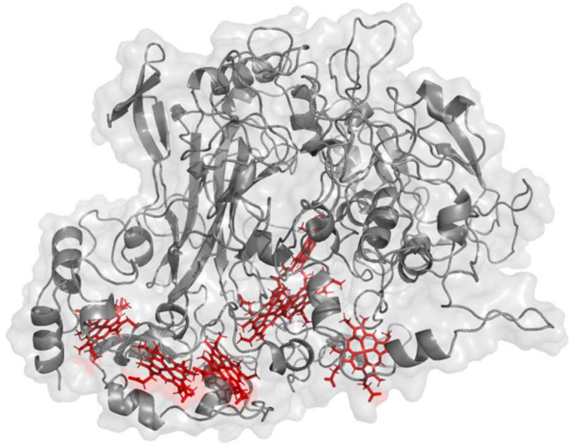

**Cluster: 13**
Sequence: WP_199384373.1
Cluster size: 28
Organism: *Geomesophilobacter sediminis*
Number of heme-binding motifs: 7
Predicted molecular weight: 102.7 kDa

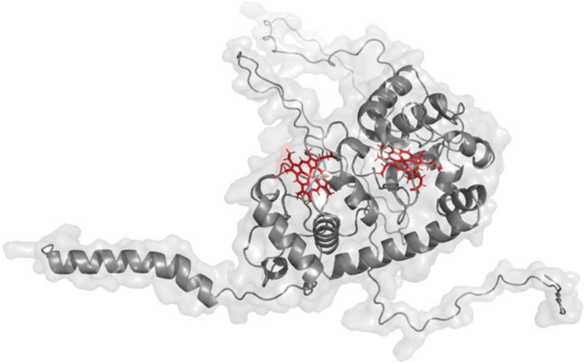

**Cluster: 14**
Sequence: WP_236025745.1
Cluster size: 28
Organism: *Geomonas azotofigens*
Number of heme-binding motifs: 2
Predicted molecular weight: 48.9 kDa

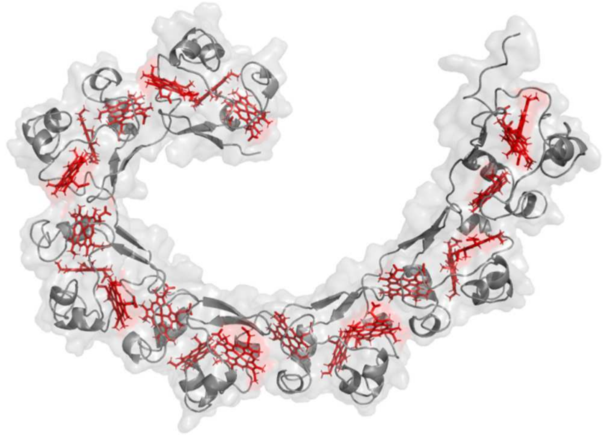

**Cluster: 15**
Sequence: WP_162148572.1
Cluster size: 26
Organism: *Desulfuromonas* sp. TF
Number of heme-binding motifs: 21
Predicted molecular weight: 70.2 kDa

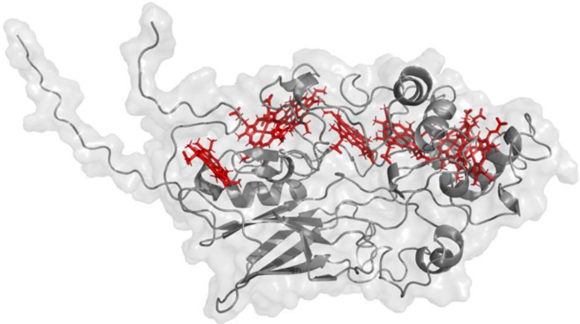

**Cluster: 16**
Sequence: Cluster size: 26
Organism: *Geobacter* sp. AOG1
Number of heme-binding motifs: 7
Predicted molecular weight: 48.4 kDa

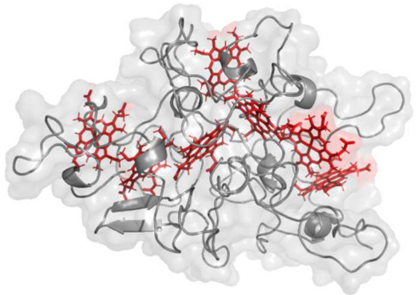

**Cluster: 17**
Sequence: WP_006001755.1
Cluster size: 23

Organism: *Desulfuromonas acetoxidans*
Number of heme-binding motifs: 7
Predicted molecular weight: 39.9 kDa

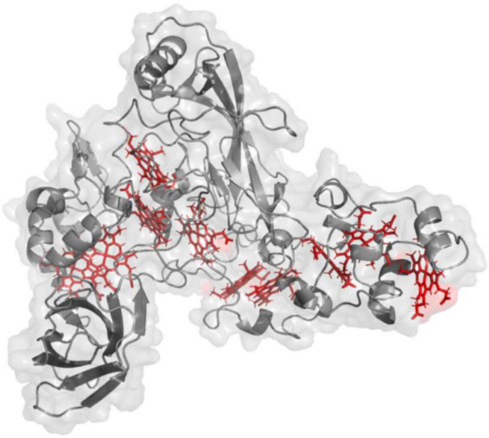

**Cluster: 18**
Sequence: WP_207163837.1
Cluster size: 21
Organism: *Geobacter benzoatilyticus*
Number of heme-binding motifs: 10
Predicted molecular weight: 68.8 kDa

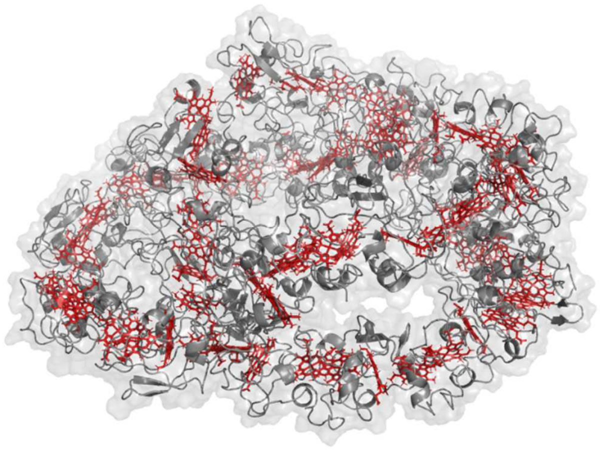

**Cluster: 19**
Sequence: WP_160295258.1
Cluster size: 19
Organism: *Geobacter* sp. OR-1
Number of heme-binding motifs: 68
Predicted molecular weight: 319.0 kDa

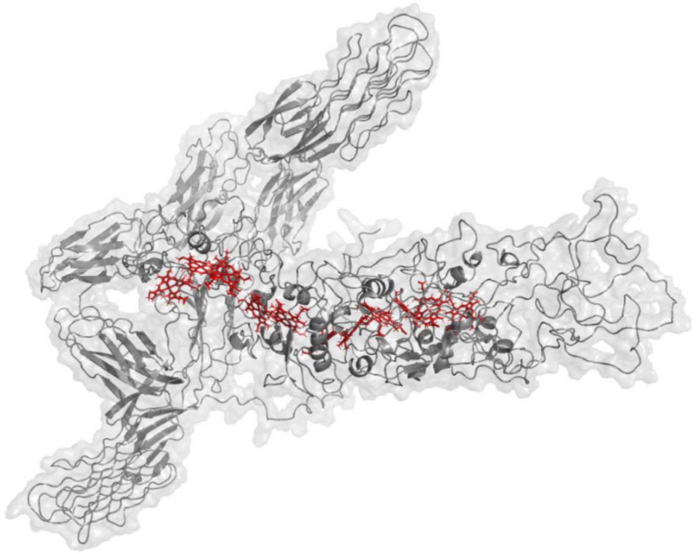

**Cluster: 20**
Sequence: WP_214174243.1
Cluster size: 18
Organism: *Geobacter luticola*
Number of heme-binding motifs: 11
Predicted molecular weight: 187.4 kDa

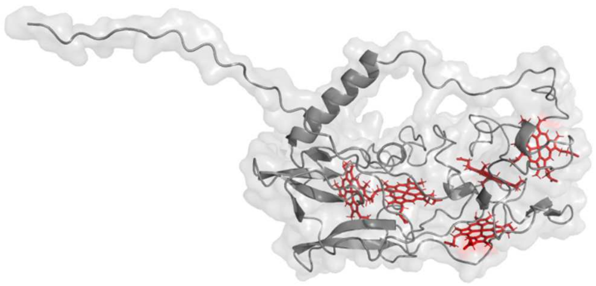

**Cluster: 21**
Sequence: WP_216499605.1
Cluster size: 18
Organism: *Geomonas diazotrophica*
Number of heme-binding motifs: 5
Predicted molecular weight: 33.9 kDa

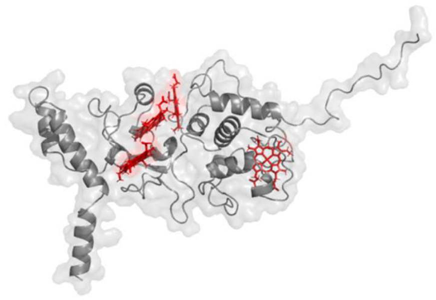

**Cluster: 22**
Sequence: WP_145025852.1
Cluster size: 17
Organism: *Geobacter argillaceus*
Number of heme-binding motifs: 4
Predicted molecular weight: 34.9 kDa

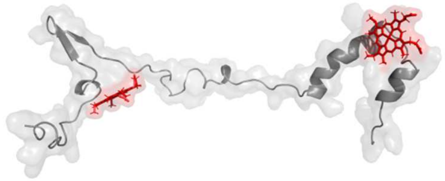

**Cluster: 23**
Sequence: WP_072283936.1
Cluster size: 17
Organism: *Syntrophotalea acetylenivorans*
Number of heme-binding motifs: 2
Predicted molecular weight: 11.3 kDa

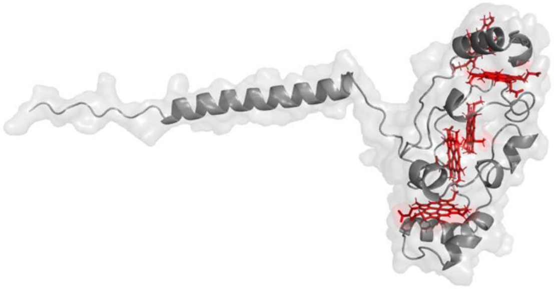

**Cluster: 24**
Sequence: WP_199384988.1
Cluster size: 16
Organism: *Geomesophilobacter sediminis*
Number of heme-binding motifs: 5
Predicted molecular weight: 23.7 kDa

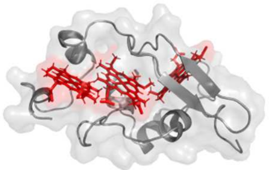

**Cluster: 25**
Sequence: WP_012774163.1
Cluster size: 16
Organism: *Geobacter* sp. M21
Number of heme-binding motifs: 3
Predicted molecular weight: 14.0 kDa

**Fig. S2.**

AlphaFold2 structure prediction of each representative cluster of MHC from which there is no available characterized reference MHC.Sequence accession NCBI code, number of sequences of the clusters, organism from which each representative sequence belongs, number of heme-binding motifs and the predicted molecular weight (hemes included) is shown bellow each predicted structure.

**Inner membrane**

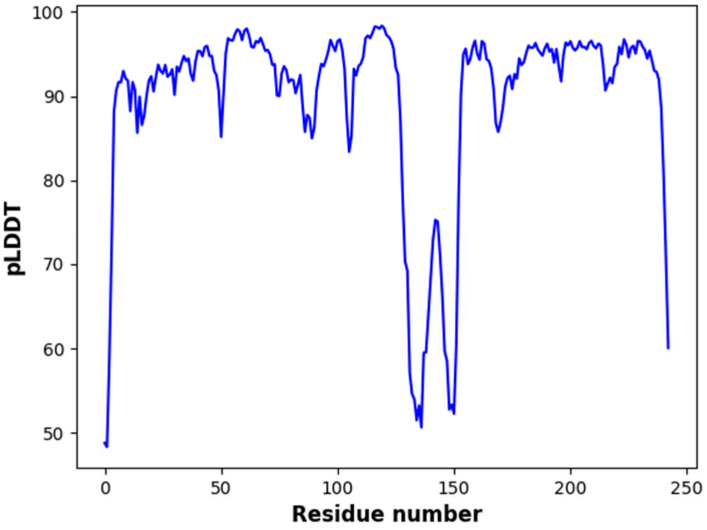


**Cluster: 1**

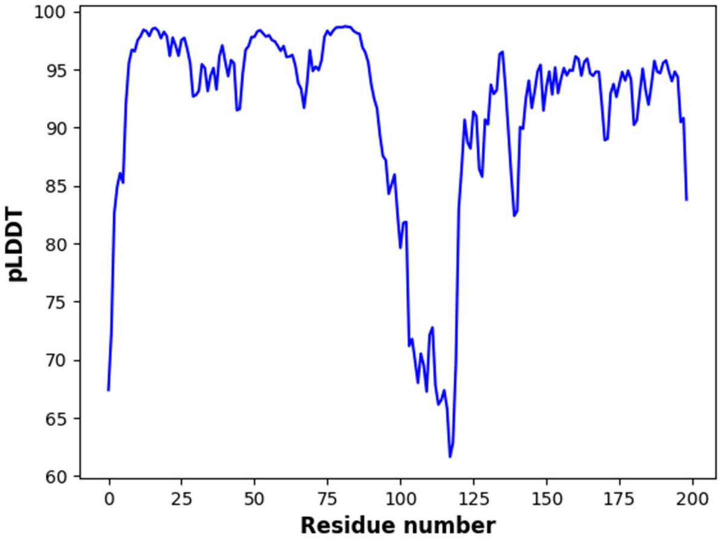


**Cluster: 2**

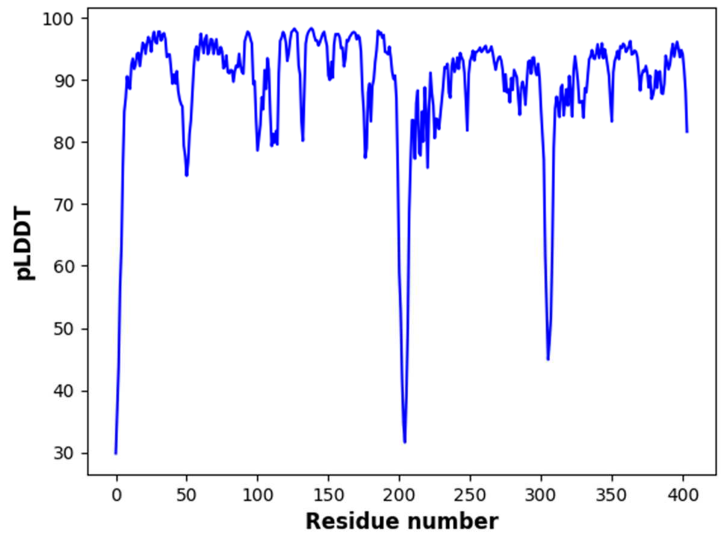

Cluster: 3

**Periplasm**

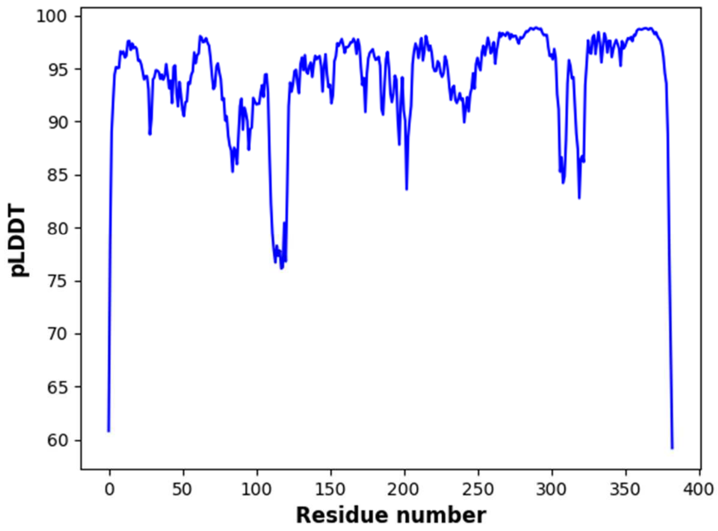

Cluster: 1

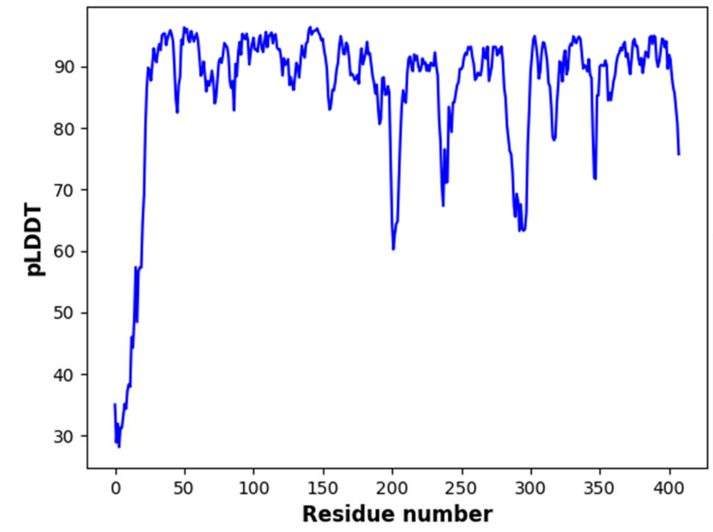


**Cluster: 2**

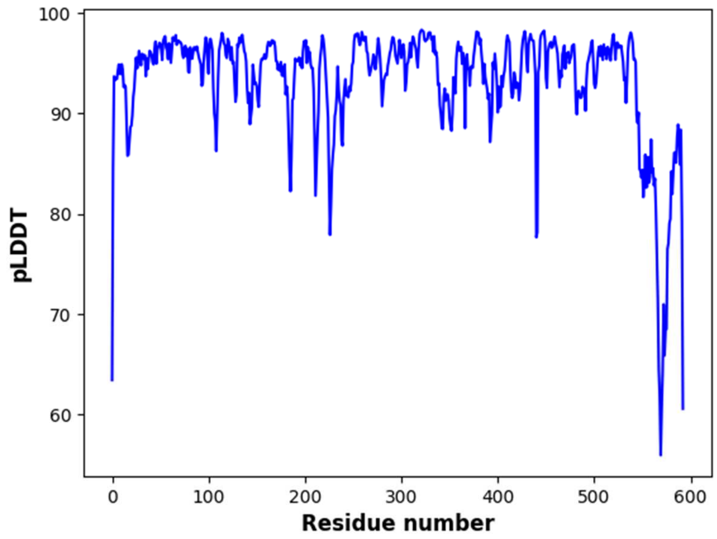


**Cluster: 3**

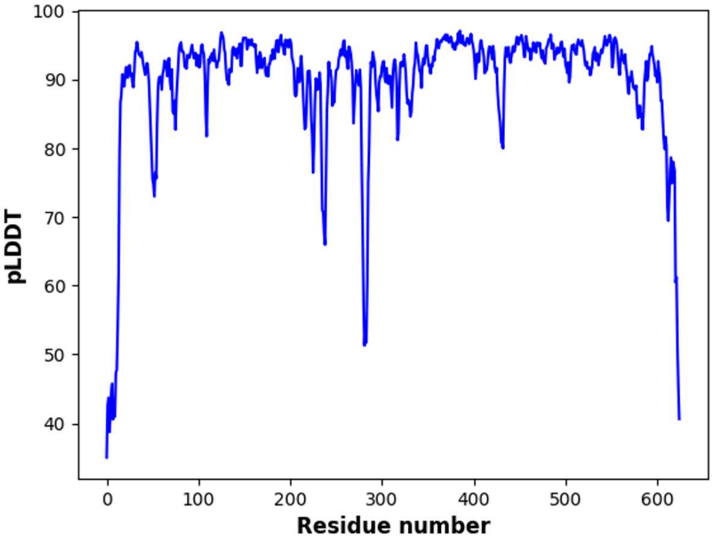


**Cluster: 4**

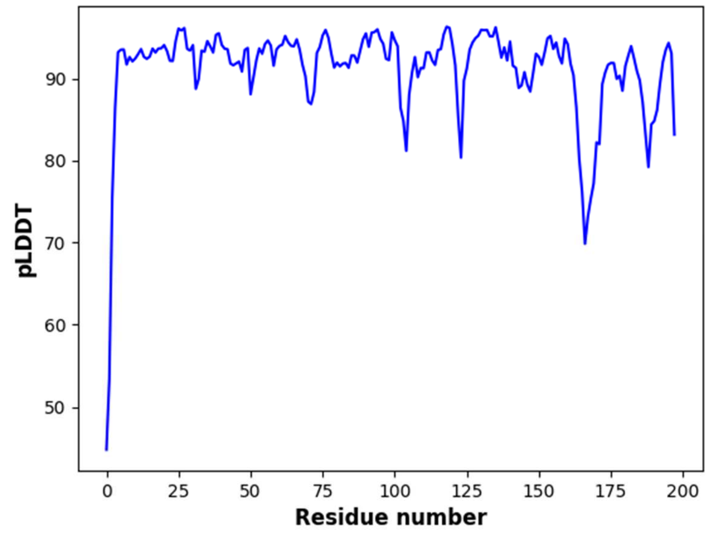


**Cluster: 5**

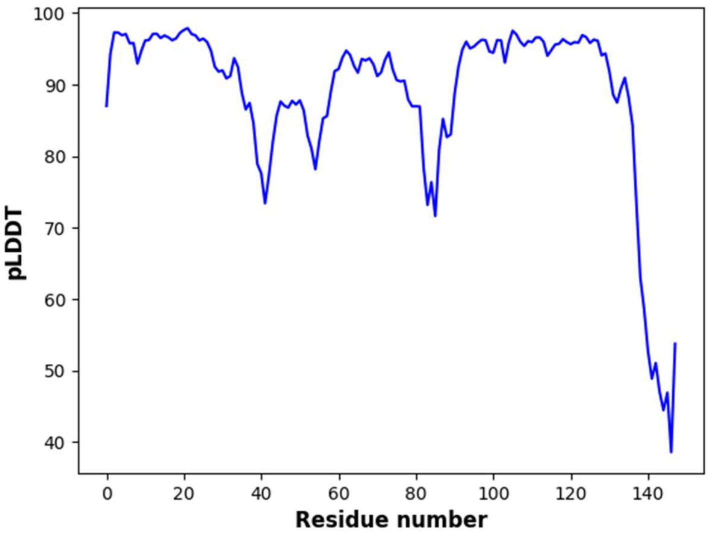


Cluster: 6

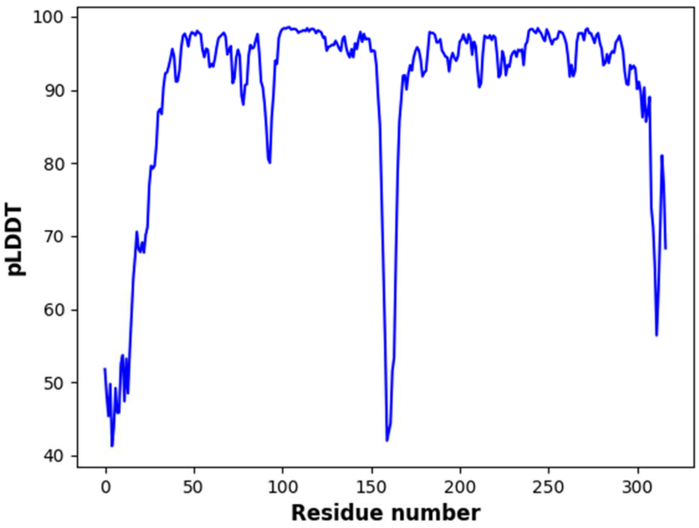


Cluster: 7

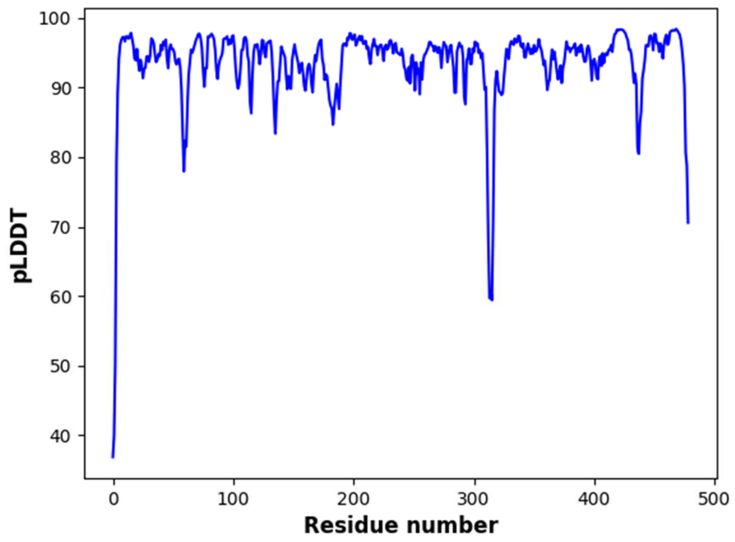


**Cluster: 8**

**Extracellular**

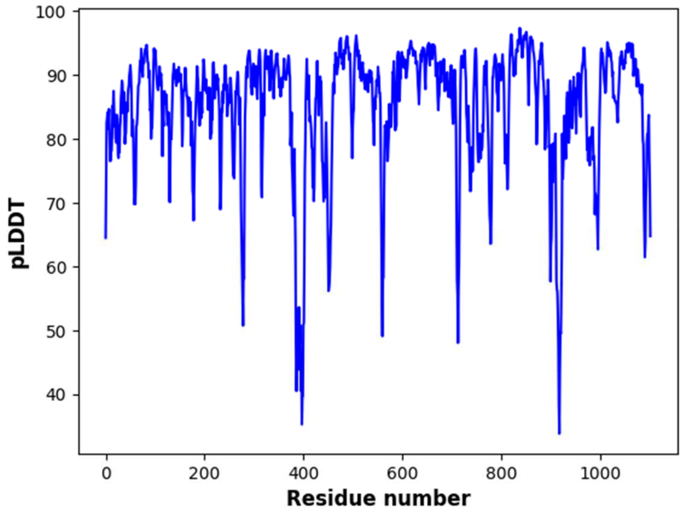


**Cluster: 1**

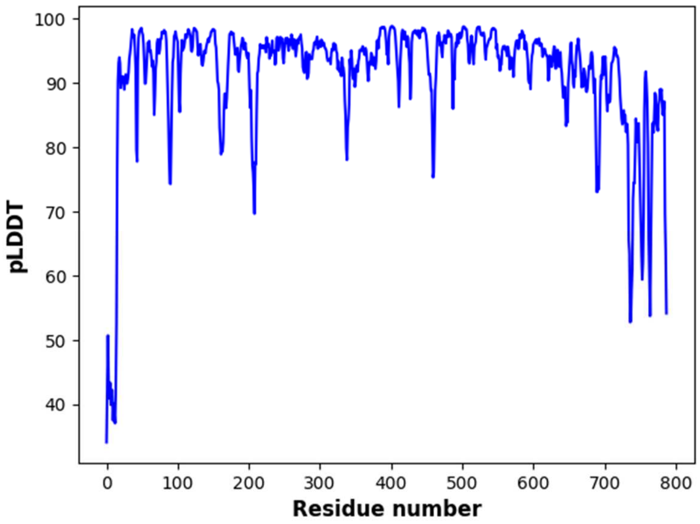


**Cluster: 2**

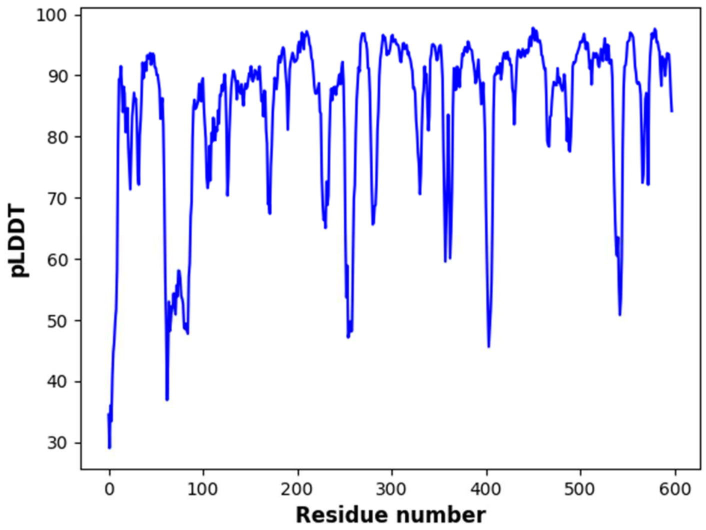


**Cluster: 3**

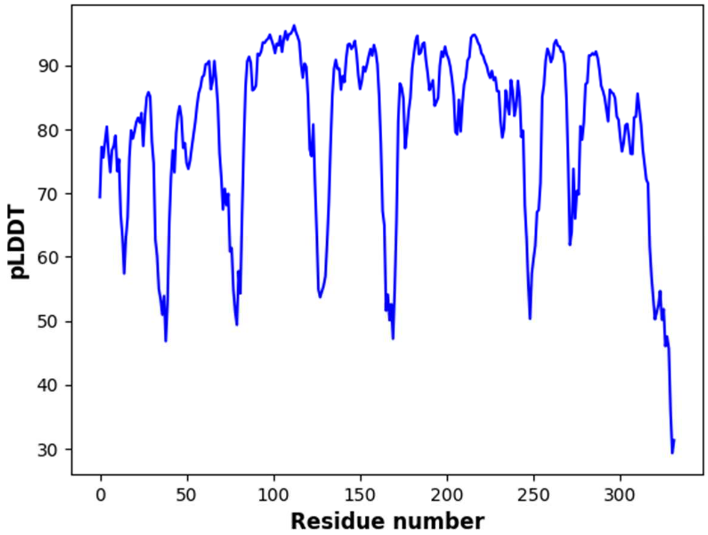


**Cluster: 4**

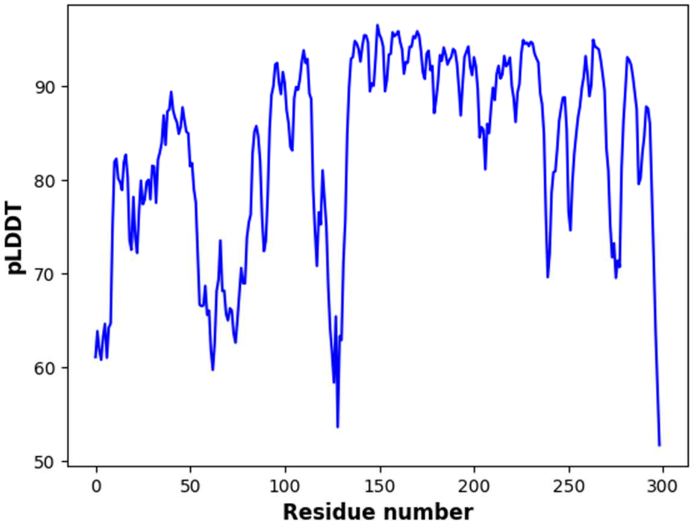


**Cluster: 5**

**Unknown location**

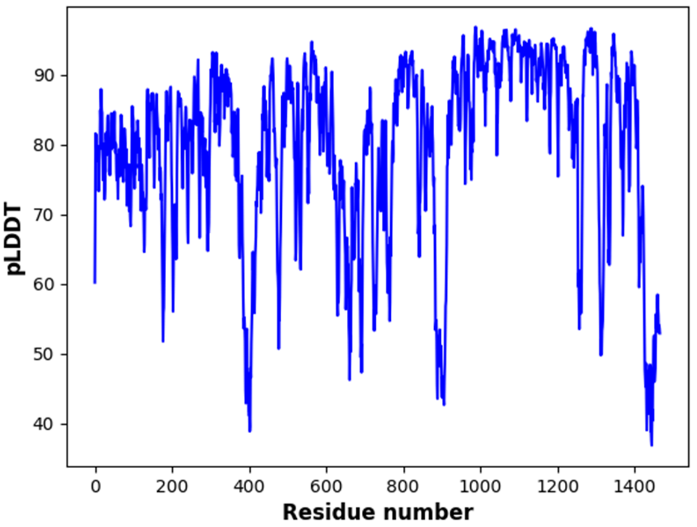


**Cluster: 1**

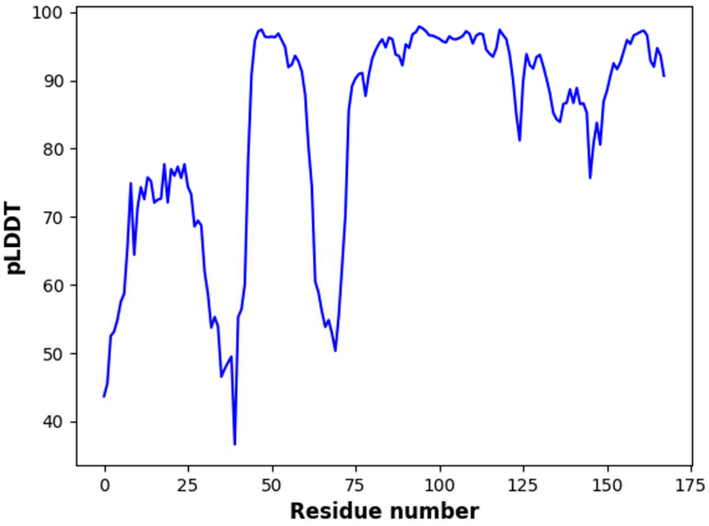


**Cluster: 2**

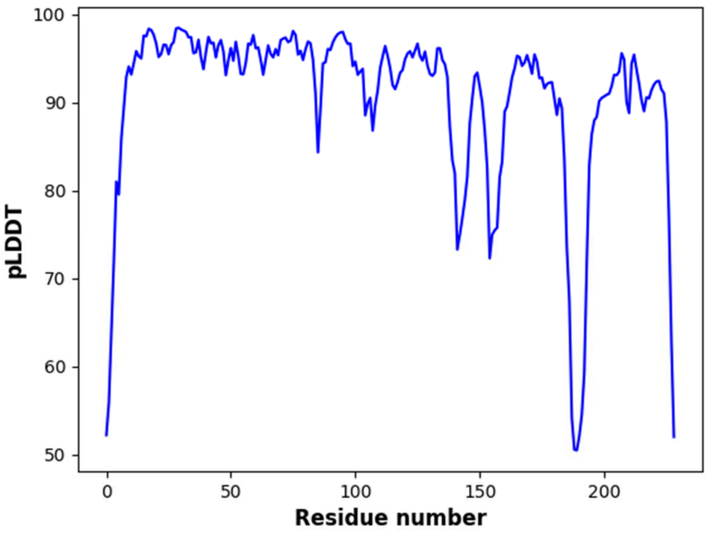


Cluster: 3

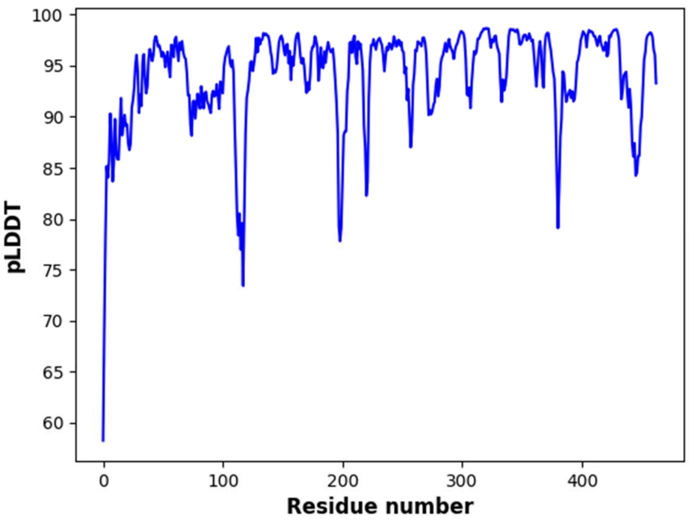


Cluster: 4

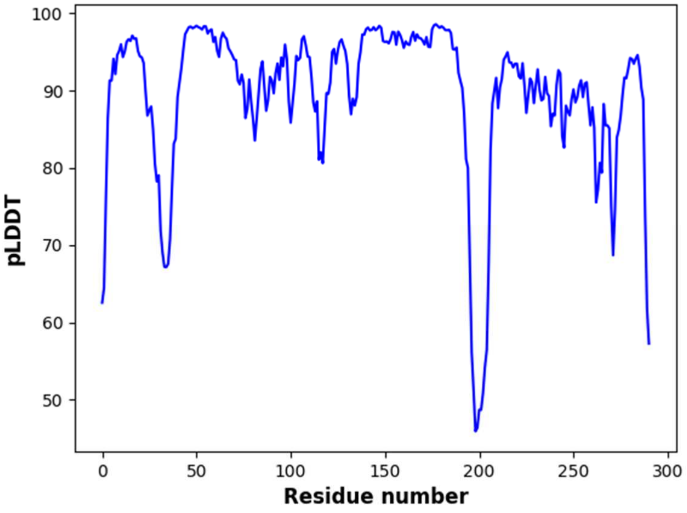


**Cluster: 5**

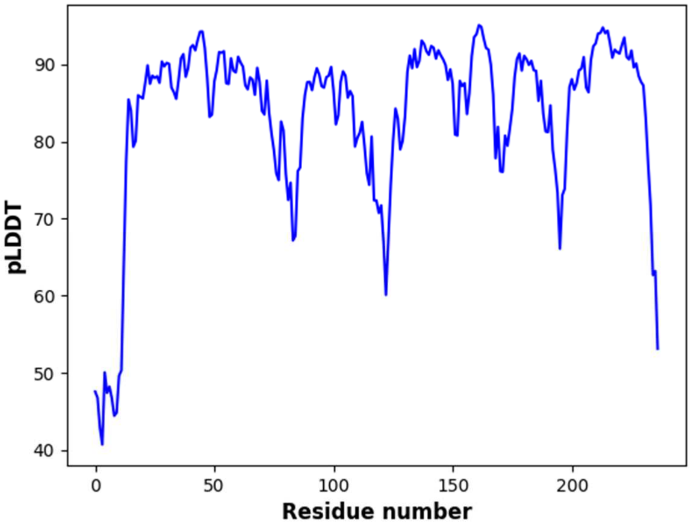


**Cluster: 6**

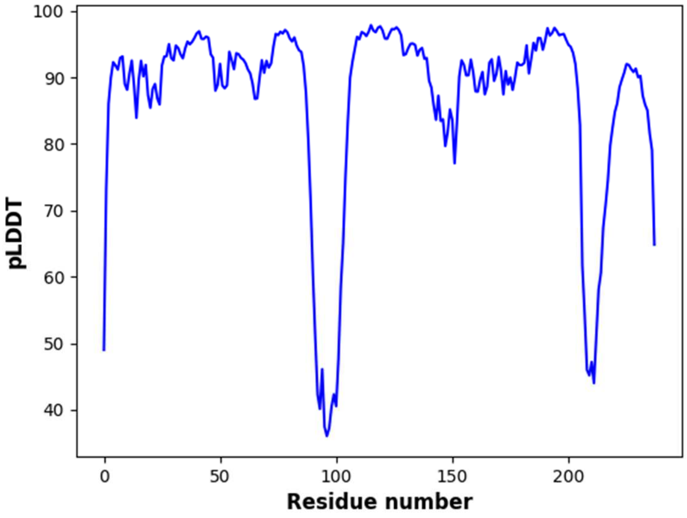

**Cluster: 7**

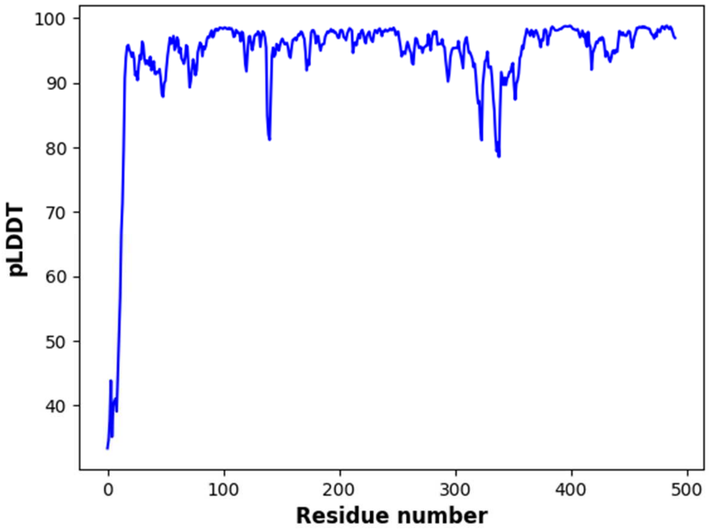

**Cluster: 8**

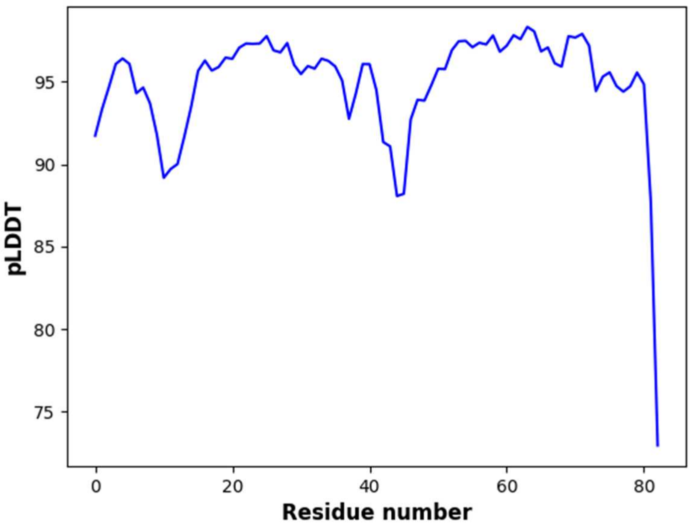


**Cluster: 9**

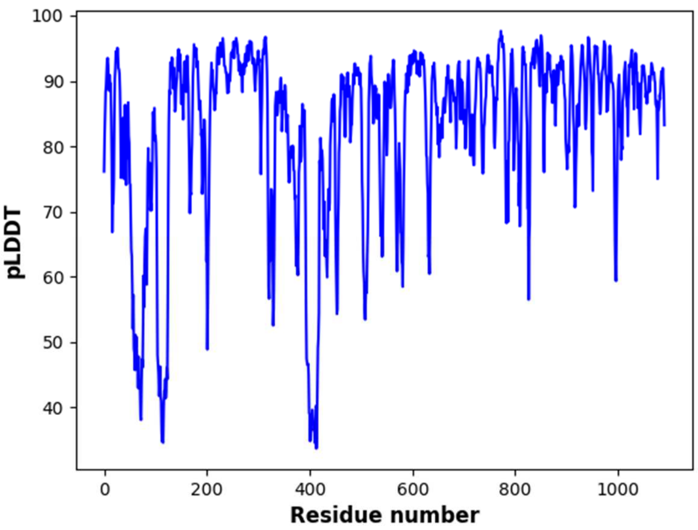


**Cluster: 10**

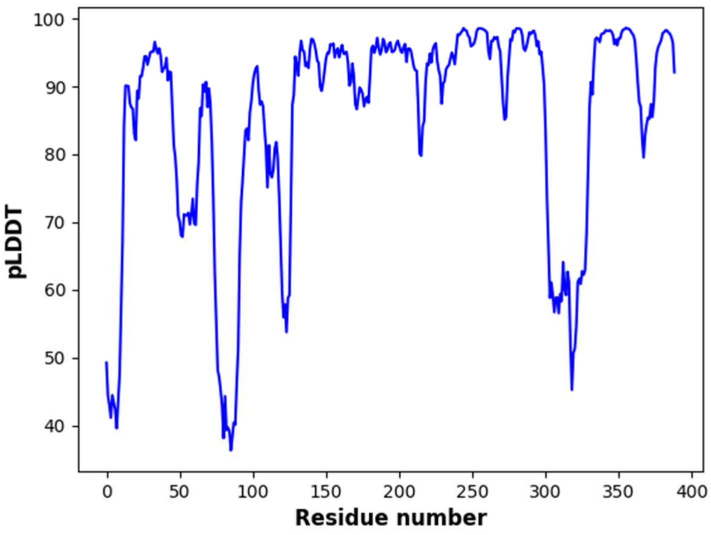


**Cluster: 11**

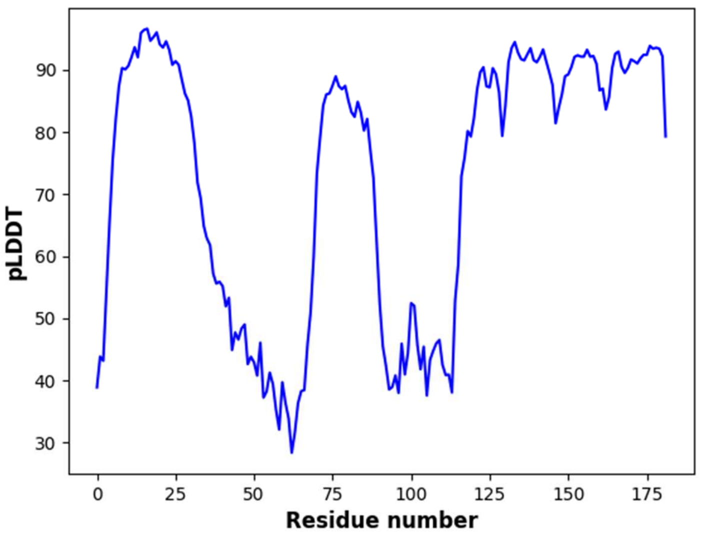


**Cluster: 12**

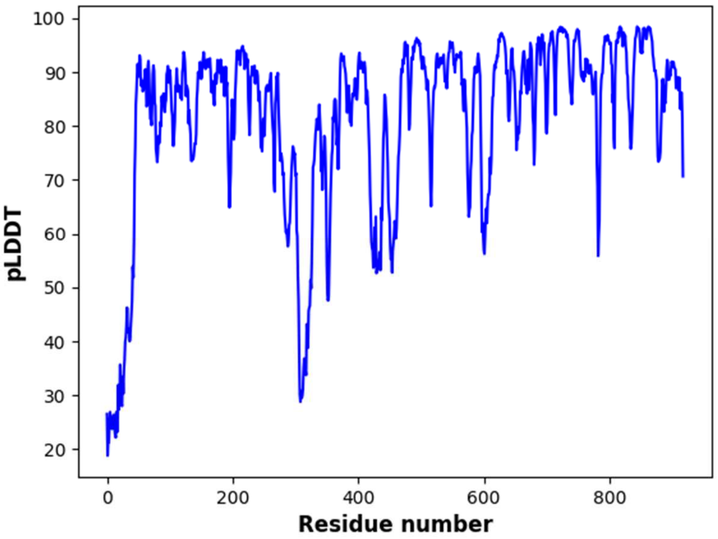


**Cluster: 13**

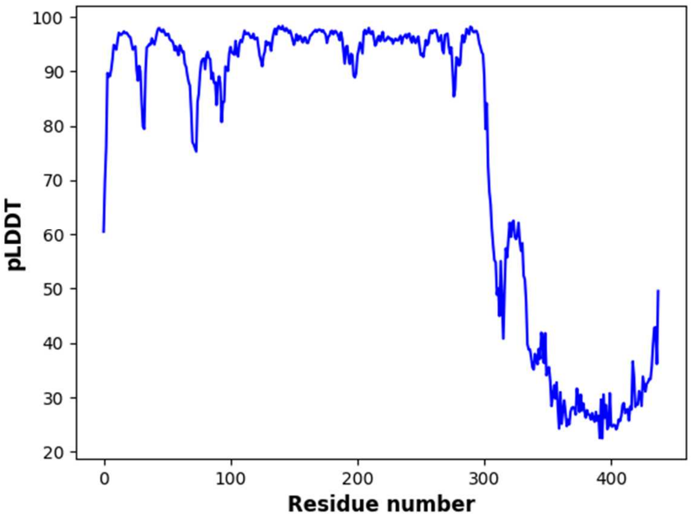


**Cluster: 14**

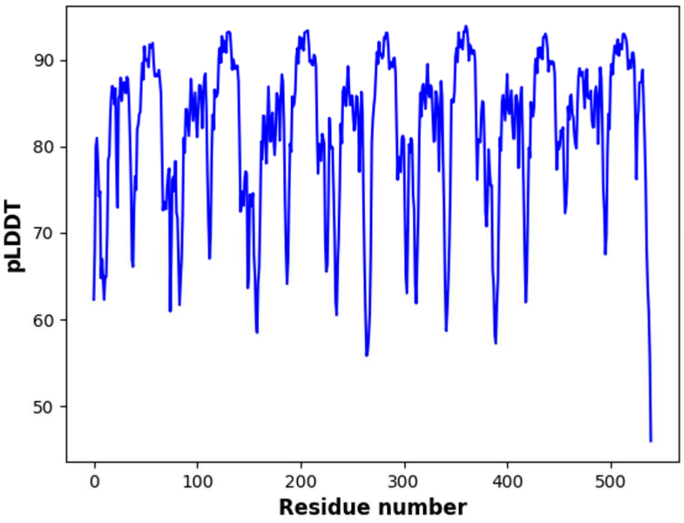

**Cluster: 15**

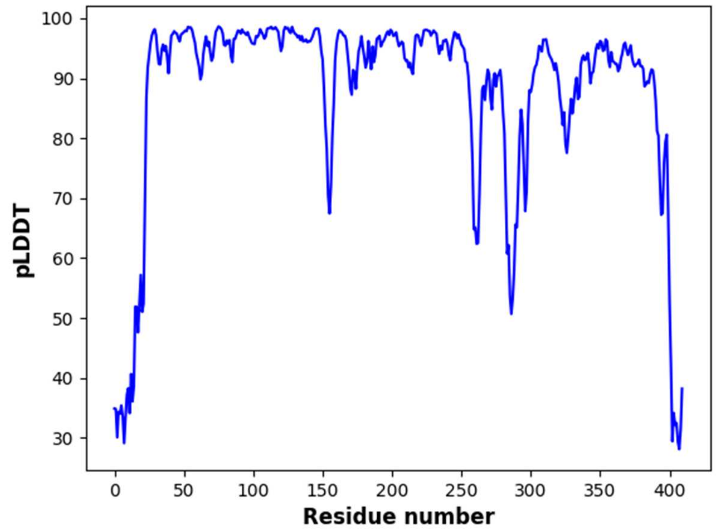

**Cluster: 16**

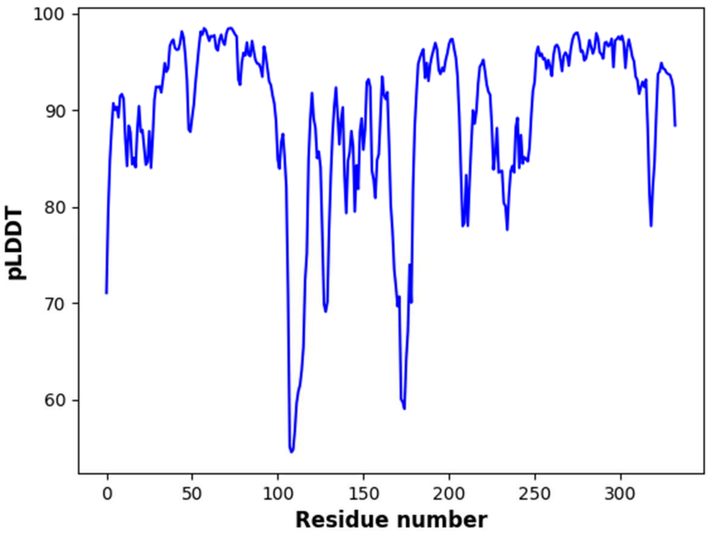


**Cluster: 17**

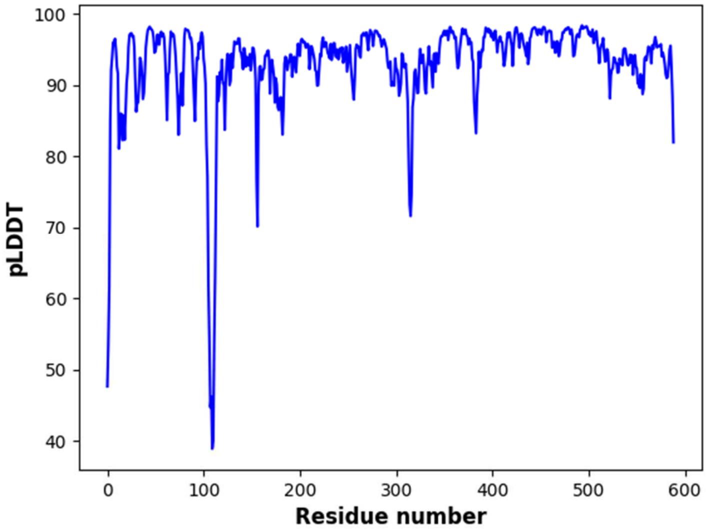


**Cluster: 18**

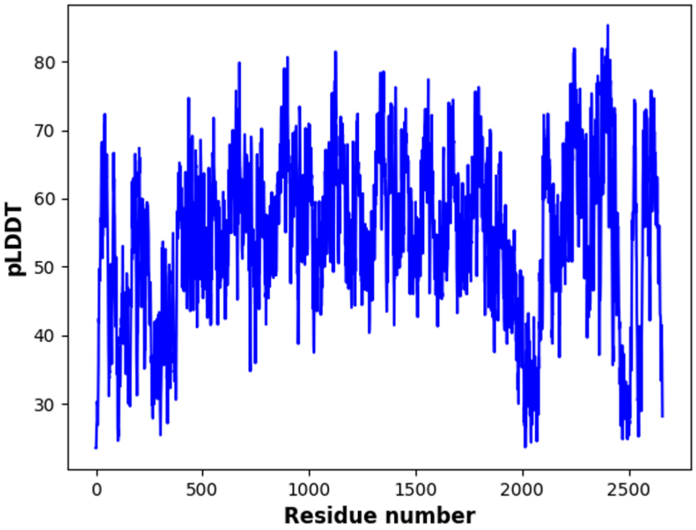


**Cluster: 19**

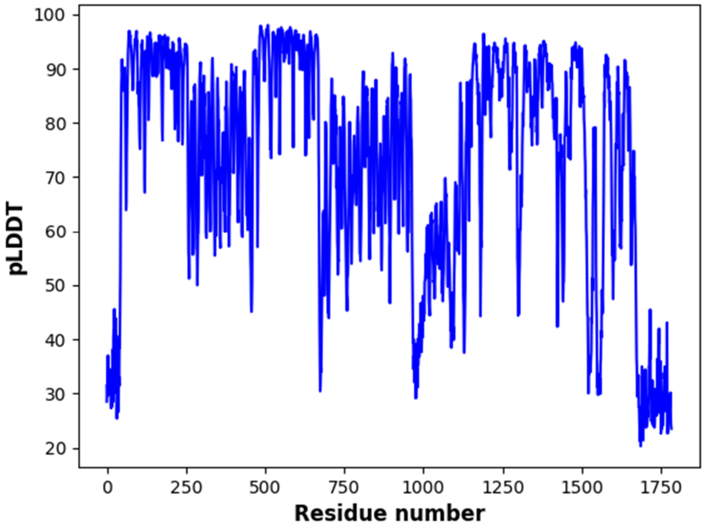


**Cluster: 20**

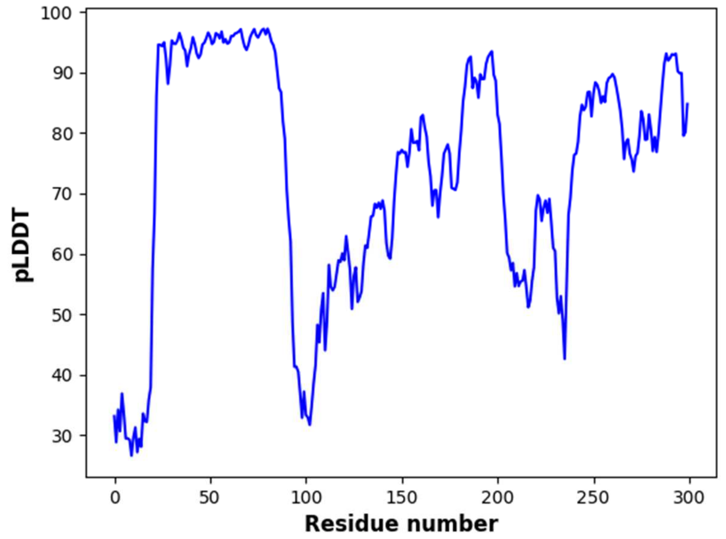


**Cluster: 21**

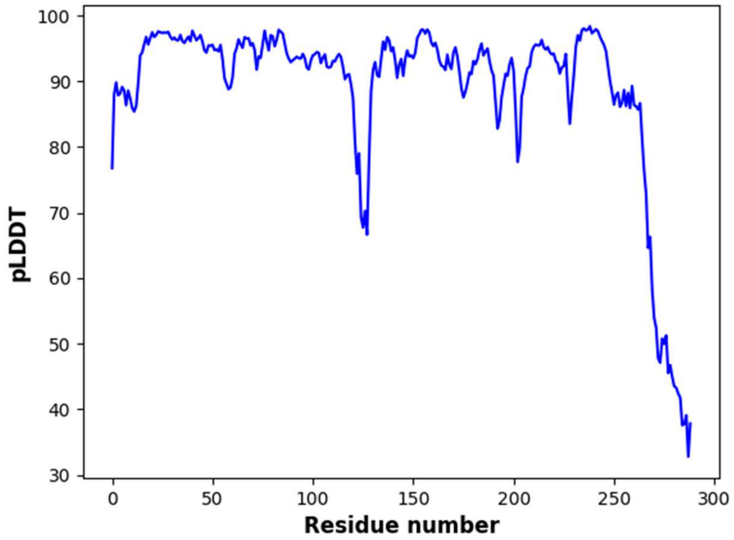


**Cluster: 22**

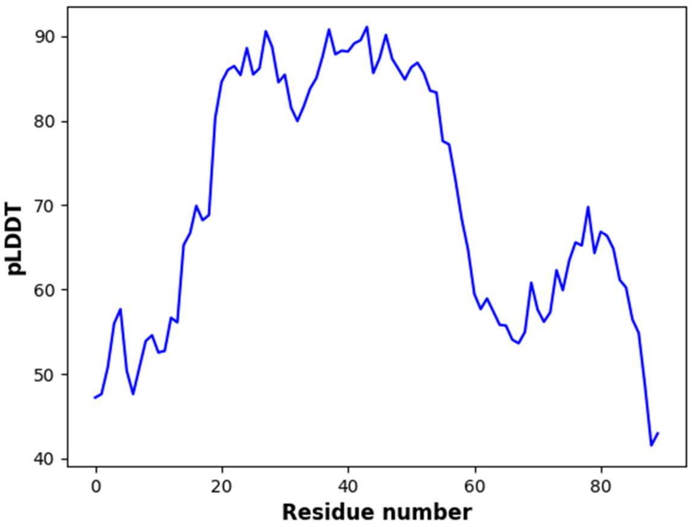


**Cluster: 23**

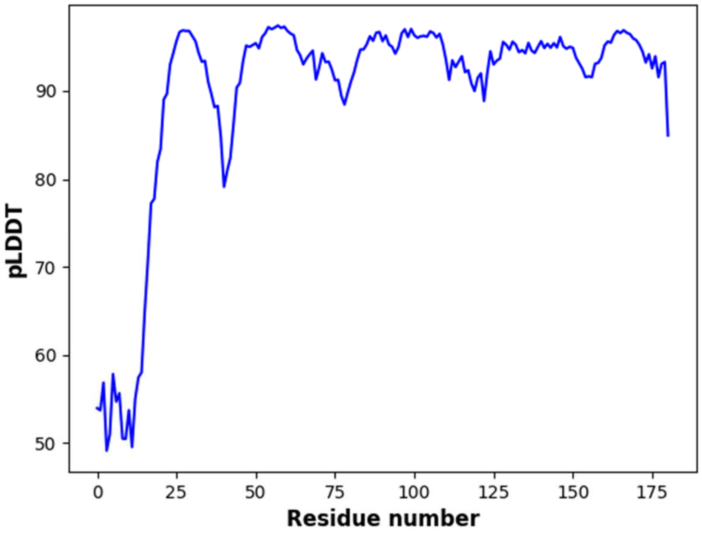


**Cluster: 24**

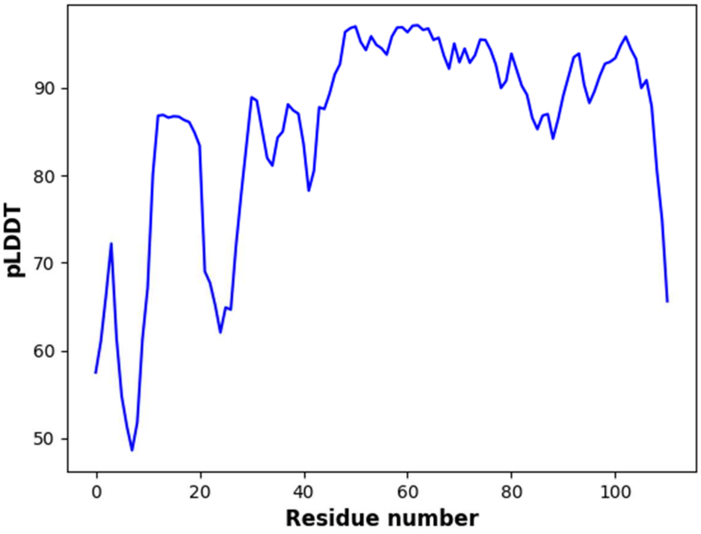


**Cluster: 25**

**Fig. S3.**

AlphaFold structure confidence prediction for each representative cluster of MHC from which there is no available characterized reference MHC (Fig. S2). Fig. S3 displays plots of the per-residue estimate of model confidence as a function of the predicted local distance difference test (pLDTT) on a scale from 0 – 100.

**Table S1.**

Number of MHC sequnces from the *Desulfuromonadia* that match each reference homologous groups of characterized MHC. MHC sequences that did not match any reference homologous groups of characterized MHC are labeled as “No identification”.

| Homologous group | Number of sequences |
|---|---:|
| No identifiation | 2147 |
| Cyt c7/PpcA/B/C/D/E | 308 |
| OmcE/OmcP | 160 |
| NrfA | 152 |
| ExtA | 145 |
| PpcF/OTR | 138 |
| OmcJ/S/T | 129 |
| GSU1996/OmcQ | 127 |
| OmaV/W | 104 |
| PccR | 102 |
| DmsE/MtoA/MtrA/OmcI | 91 |
| BthA/MacA/MauG/PsaCCP | 87 |
| DHC2 | 76 |
| CbcL | 74 |
| OmaC/B | 73 |
| CbcA | 70 |
| ImcH | 65 |
| CbcS | 58 |
| OmcH/O | 57 |
| CbcX | 57 |
| CymA/NapC/NrfH | 56 |
| ExtK | 49 |
| OmcB/C | 43 |
| PccG | 40 |
| FccA | 38 |
| GSU0616 | 37 |
| GSU0105 | 34 |
| STC | 27 |
| GSU1334/OmcZ | 26 |
| ActA | 22 |
| GSU1786 | 20 |
| OmcG | 17 |
| GSU2884(OmcA) | 17 |
| IhOCC | 14 |
| ExtD | 12 |
| ExtG | 8 |
| Split/Soret cyt c | 8 |
| PgcA | 7 |

| | |
|---|---|
| OmcM | 7 |
| HAO/HDH | 5 |
| Cyt c3 | 2 |
| MtrC/F | 2 |
| DsrJ | 2 |
| QHNDH | 1 |
| MccA | 1 |
| sb-DHC | 1 |

**Data S1.(separate file)**

General characteristics of the genomes collected from the NCBI RefSeq database.

**Data S2.(separate file)**

Full dataset of the MHC protein sequences from the *Desulfuromonadia* in fasta format.

**Data S3.(separate file)**

Database of characterized MHC used to identify the full dataset of the *Desulfuromonadia* MHC. Sequences are in fasta format with the header contains the protein name, flowed by the organism that it belongs and the NCBI accession code.

**Data S4.(separate file)**

Identification results for the of the full dataset of MHC of the *Desulfuromonadia*

**Data S5.(separate file)**

Structure files in PDB format for the structures of represented in Fig. S1 and S2. Structures of the characterized reference MHC contain the AlphaFold database accession code within each file name
